# Gangliosides GM3 And GD3 Modulate Insulin Aggregation Pathways and Reduce Cytotoxicity Through Structural Remodeling

**DOI:** 10.64898/2026.02.03.703542

**Authors:** Nazifa Tasnim Ahmad, Jhinuk Saha, Yimin Mao, Robert Silvers, Zaid Abulaban, Joshua Mysona, Ayyalusamy Ramamoorthy

**Affiliations:** Department of Chemical and Biomedical Engineering, FAMU-FSU College of Engineering, 2525 Pottsdamer St., Tallahassee, FL 32310, United States; National High Magnetic Field Laboratory, 1800 E. Paul Dirac Drive, Tallahassee, FL 32310, United States; Department of Chemistry and Biochemistry, Florida State University, Tallahassee, FL, USA; Institute of Molecular Biophysics, Florida State University, Tallahassee, FL 32304, United States

**Keywords:** Insulin aggregation, Gangliosides, Type-2 Diabetes (T2D), Polymorphism, Cytotoxicity, Non-fibrillar aggregates

## Abstract

Insulin amyloid aggregation is a key pathological and pharmaceutical concern, particularly in the context of Type-2 Diabetes (T2D), where amyloid deposition of protein can impair therapeutic efficacy and contribute to cell death leading to local tissue damage. Although gangliosides—glycosphingolipids containing sialic acid residues—are known to modulate amyloid formation in neurodegenerative disorders, their influence on insulin aggregation remains largely unexplored. In this study, we investigate the effects of gangliosides GM3 and GD3 on insulin aggregation. Using Thioflavin-T (ThT) based fluorescence kinetics, Fourier Transform Infrared (FTIR) spectroscopy, Circular Dichroism (CD) spectroscopy, Small Angle X-ray Scattering (SAXS), Nuclear Magnetic Resonance (NMR) spectroscopy, and Transmission Electron Microscopy (TEM), the aggregation pathway, changes in the secondary structure and morphology of insulin aggregates have been characterized. Our results show that both GM3 and GD3 lipids accelerated insulin aggregation in a concentration-dependent manner while steering the pathway away from classical fibril formation, producing short, beaded structures distinct from the extended fibrils observed under lipid-free conditions. CD and FTIR data analyses revealed that insulin in the presence of gangliosides formed non-fibrillar intermediates with distinct secondary structures: β-sheet-rich globular clusters in presence of GD3 and α-helical intermediates in GM3-treated samples. Cytotoxicity assays further demonstrated that ganglioside-induced aggregates are significantly less toxic to cells when compared to insulin-only aggregates. Furthermore, ganglioside-bound insulin oligomers retain seeding capacity, suggesting that they can nucleate further aggregation despite their non-fibrillar morphology. These findings underscore the role of gangliosides in modulating insulin amyloid polymorphism and toxicity, offering new insights into their potential impact on the pathology of T2D and treatment strategies.

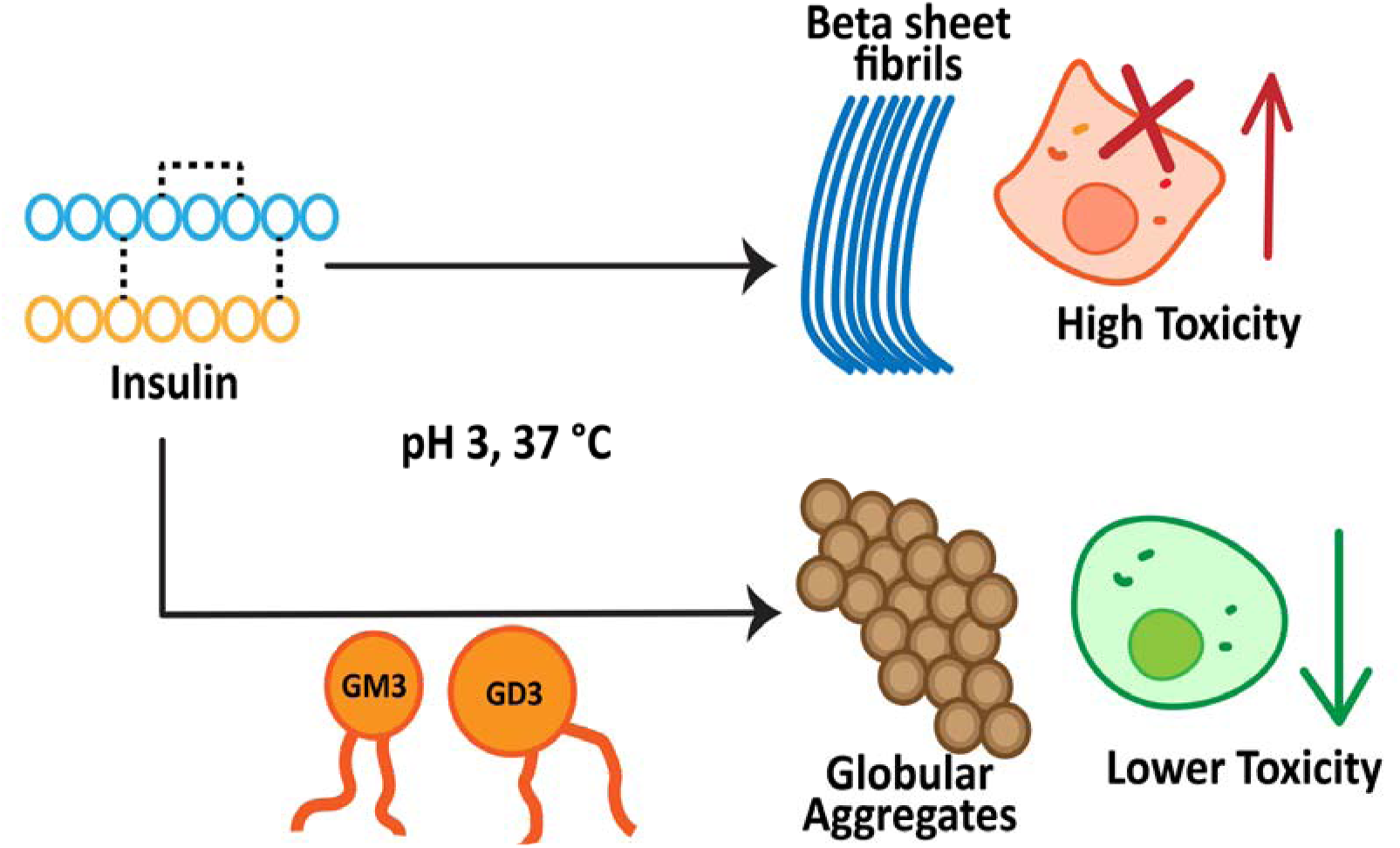

**Highlights:**

- Gangliosides GD3 and GM3 accelerate insulin aggregation, forming non-fibrillar assemblies.
- Ganglioside-bound insulin aggregates are less cytotoxic than fibrillar aggregates.
- Despite altered morphology, ganglioside-bound aggregates retain seeding ability.

## 1. Introduction

Insulin is a hormone containing 51 amino acids in two peptide chains, A and B, connected by two interchain and one intrachain disulfide bonds [1,2]. It is synthesized as preproinsulin and undergoes proteolytic processing in endoplasmic reticulum and Golgi bodies in the pancreatic β-cells to emerge as the active monomeric form, which is then stored in a zinc-bound hexameric form in secretory granules [2]. Its primary role is to maintain glucose homeostasis in blood by facilitating glucose uptake in cells, glycogenesis and gluconeogenesis [3,4]. Beyond its central role in glucose homeostasis, insulin is also an amyloidogenic protein due to its inherent propensity to misfold and aggregate under destabilizing conditions. Depending on the local environment, insulin monomers can undergo unfolding to form amyloid fibrils characterized by peptide strands organized cross β-sheet structures stabilized by hydrogen bonds [5–8].

Protein amyloidogenesis is a pathological hallmark of various diseases, including Alzheimer’s disease, Parkinson’s disease and Type 2 Diabetes (T2D). These involve deposition of amyloid aggregates of proteins in tissue within their pathophysiology [9–11]. In the context of diabetes, insulin aggregation is particularly relevant. In type I and II diabetes, where exogeneous insulin administration is essential [12], chronic subcutaneous injections can lead to the formation of localized amyloid deposits, termed insulin-derived amyloidosis or insulin amyloidoma [13–20]. These deposits are unavailable for cellular uptake and metabolism, and they can cause obstruction in insulin absorption and impair glycemic control. This can lead to the loss of therapeutic efficacy, local inflammation and cytotoxicity to the surrounding tissues. Several patients have been observed to develop tissue necrosis and severe infection around the injection sites as well [17,21–23]. Additionally, the need for higher insulin doses in patients with amyloid-related complications increases the expense to patients, further adding to the growing economic burden of the disease [24]. The process of amyloid formation may be accelerated by various kinds of stress, including elevated temperature, extreme pH and salinity, the presence of small biomolecules, surfactants and organic solvents, and mechanical force[5,8,25–28]. Among these various modulators, lipids have emerged as key regulators of amyloidogenesis[29].

Lipids and cell membranes have been shown not only to catalyze protein aggregation but also to significantly influence the kinetics and pathways of aggregation, as well as the structural characteristics and toxicity of the resulting amyloid species. These effects are highly dependent on the lipid composition and factors such as lipid head group charge, acyl chain saturation level, which can alter membrane properties and protein-lipid interactions. In subcutaneous fat, injected insulin interacts with various lipids, including phospholipids, ceramides, and sphingolipids present in cell membranes [30]. Matveyenka and coworkers studied the effect of these lipids and their mixtures on insulin aggregation [30]. They found that anionic phospholipids such as cardiolipin (CL), phosphatidylserine (PS), and ceramide (CER) significantly shortened the lag-phase and accelerated insulin aggregation. Whereas sphingomyelin (SM) and Zwitterionic Phosphatidylcholine (PC) delayed and inhibited insulin aggregation, respectively. While previous studies on lipid–insulin interactions have focused on anionic or zwitterionic phospholipids, these models may not completely reflect physiologically relevant conditions. Anionic phospholipids (such as PS) are largely localized to the cytosol-facing inner leaflet of the plasma membrane [31–35], whereas insulin is synthesized in pancreatic β-cells and secreted into the extracellular space via secretory vesicles [36]. This spatial disconnect suggests that insulin is more likely to interact with lipids in the outer leaflet of the plasma membrane rather than those on the cytosolic side. Among the lipids of the outer membrane, gangliosides are one of the most prominent anionic species. Gangliosides are glycosphingolipids composed of a ceramide tail and an oligosaccharide headgroup that contains one or more sialic acid residues [37]. They account for up to 15 mol% of outer leaflet lipids in red blood cells and are known to cluster into lipid rafts in cholesterol-rich membranes [32,38,39]. These contain negatively charged domains that can interact with positively charged proteins like insulin, facilitating localized protein–lipid interactions. Gangliosides have been considerably implicated in amyloid aggregation associated with neurodegenerative diseases, most notably Alzheimer’s and Parkinson’s disease [40,41]. In Alzheimer’s pathology, GM1 ganglioside clusters on the membrane surface have been shown to act as nucleation centers for amyloid-β (Aβ) aggregation [42]. Upon binding to GM1-rich lipid rafts, Aβ undergoes conformational changes from α-helix to β-sheet, accelerating fibril formation [43–46]. The resulting GM1-bound Aβ complexes (GAβ) have been identified in early AD brain tissue and are thought to serve as seeds for further aggregation [47,48]. Despite their well-established role in modulating amyloid aggregation of Aβ, the influence of gangliosides on insulin aggregation remains largely unexplored. Given the spatial proximity of gangliosides and insulin in the extracellular space, understanding how these lipids affect insulin’s structural fate may yield important insights into insulin-derived amyloid pathology. In this study, we investigated the effects of gangliosides GD3 (Fig. S1.a) and GM3 (Fig. S1.b) on human insulin aggregation. Employing ThT fluorescence-based kinetics assay, FTIR spectroscopy and CD spectroscopy, TEM and cytotoxicity assays, we measured how GM3 and GD3 modulate the aggregation pathway, morphology, secondary structure and toxicity of insulin aggregates. Our findings reveal that both GM3 and GD3 accelerate insulin aggregation but divert it from the classical fibril-forming pathway towards a non-fibrillar off-pathway oligomers with distinct secondary structure and markedly reduced cellular toxicity.

## 2. Experimental Section

### 2.1. Materials

Recombinant human insulin was purchased from Roche, USA (Indianapolis, IN). Monosialodihexosylganglioside GM3 and Disialotetrahexosylganglioside GD3 lipids were acquired from Avanti Polar Lipids (Alabaster, Alabama). Sodium phosphate (NaH_2_PO_4_), sodium chloride (NaCl) and Trypsin-EDTA were obtained from Fisher Scientific (Hampton, NH). Tris (2-carboxyethyl) phosphine hydrochloride (TCEP HCl) was obtained from Sigma-Aldrich (St. Louis, MO, USA). Copper grids for transmission electron microscopy were supplied by Millipore Sigma (Burlington, MA), and Uranyless Stain was purchased from Electron Microscopy Sciences (Hatfield, PA, USA). Thioflavin T (ThT) dye was obtained from Millipore Sigma (Burlington, MA). Dulbecco’s Modified Eagle Medium (DMEM) was procured from Gibco (Grand Island, NY).

### 2.2. Peptide and Lipid Samples Preparation

Human Recombinant Insulin was dissolved in pH 3 sodium phosphate buffer (10 mM) while the reduced insulin formation was induced by incubating 2 mg/mL monomer solution with 50 mM TCEP HCl for an hour at room temperature. The final concentrations of the insulin solutions were measured using NanoPhotoMeter NP80 (IMPLEN) with extinction coefficient of 3360. Ganglioside stock solutions were prepared in a 1:1 mixture of chloroform and methanol at a concentration of 2.5 mg/mL. These stock solutions were stored at -20 °C until use. Lipids were then aliquoted from the stock solutions before use, dried to a film under a stream of air and then further dried under a vacuum overnight. The resulting films were subsequently resuspended in pH 3 phosphate buffer for one hour.

### 2.3. Thioflavin T Fluorescence Assay

The kinetics of protein aggregation were measured using a Thioflavin-T (ThT)-based fluorescence assay, carried out on a Biotek Synergy H1 instrument with excitation at 452 nm and emission at 485 nm. All samples for ThT assay were prepared with 80 μM insulin in a sodium phosphate buffer (10 mM, 150 mM NaCl, pH 3), with 10 μM ThT and lipid concentrations as noted in figures. Insulin was dissolved directly in the same phosphate buffer and added to the samples immediately prior to the start of the measurement. Samples for measuring the kinetics of reduced-insulin aggregation were prepared in a similar manner. A volume of 50 μL of each sample was added to each well in a black walled 384-well plate with an opaque, flat (Corning, Ref# 3575) in triplicates. Fluorescence was measured under shaking at 700 rpm with an interval of 15 minutes at 37 °C. All ThT assays were independently replicated at least twice to ensure reproducibility. The ThT fluorescence kinetics data were analyzed in MATLAB (MathWorks, R2023a) by fitting the normalized data to a sigmoidal growth equation using nonlinear least squares. From the fitted curves, the lag time (t_lag_), half time (t□/□) and maximum fluorescence intensity (f_max_) were obtained to quantify the nucleation and growth phases of insulin aggregation.

### 2.4. Transmission Electron Microscopy

Samples for transmission electron microscopy were freshly prepared by applying 10 μL of each sample to 200 mesh-size Formvar/Carbon supported copper grids (Electron Microscopy Sciences, catalog number: FCF200-Cu-50) and dried for 10 minutes at room temperature. Then, the grids were stained with 10 μL of Uranyless stain (Electron Microscopy Sciences Catalog #22409). Excess stain solution was soaked on bloating paper, and the grid was dried for 15 more minutes at room temperature. Samples were imaged using HT7800 Hitachi TEM microscope at an acceleration voltage of 100 kV. Images were obtained from at least three grid regions for independent samples at magnifications between 15,000×-30,000×.

### 2.5. Circular Dichroism Spectroscopy

After 42 hours of incubation, all samples were diluted with sodium phosphate buffer to achieve a final insulin concentration of 20 μM. CD spectra between 200 - 280 nm were obtained using a Chirascan instrument at room temperature. The average of three scans for each sample was plotted using OriginPro software. CD spectra within the 200–250 nm range were analyzed using the BeStSel software (https://bestsel.elte.hu/) to estimate secondary structure content [49]. Deconvolution was performed by uploading baseline-corrected spectra in millidegrees (mdeg), with a specified pathlength of 0.1 cm and a protein concentration of 80 μM. The output provided the estimated percentages of α-helix, parallel and antiparallel β-sheet structures, turns, and unordered structures, based on a reference database optimized for β-structure topology.

### 2.6. Fourier Transform Infrared Spectroscopy

A total of 0.3 mg of lyophilized insulin samples was used to acquire IR spectra in the range of 1500-1800 cm^-1^ at a resolution of 8 cm^-1^ on a Thermo Nicolet instrument equipped with a diamond-ATR accessory. For example, 512 spectral scans were accumulated to enhance the signal-to-noise ratio. The resulting spectra were baseline-corrected and plotted using OriginPro Software. Peak fitting was performed using a combination of Gaussian and Lorentzian functions, with initial peak positions guided by second derivative analysis to identify spectral components corresponding to β-sheet, α-helix, turn and random-coil structures. The area under each fitted peak was calculated to estimate the relative proportions of these secondary structural elements.

### 2.7. NMR Spectroscopy

1D ^1^H NMR spectra were recorded using the *zgesgppe* pulse program and water suppression using excitation sculpting with gradients [50,51] at 298 K on a Bruker AVANCE III NMR spectrometer operating at 700 MHz equipped with a TCI cryoprobe and operating TopSpin 3.6.5. NMR spectra under low-salt conditions were recorded on samples containing 80 μM insulin in the absence and presence of 1x and 5x molar ratio GM3 in a buffer containing 10 mM NaPi pH 3.0, 10% D_2_O. Spectra were acquired with 32,768 points, 128 scans, 15.94 ppm spectral width, 3290.70 Hz transmitter frequency offset, and a recycle delay of 1.0 s. NMR spectra under salty conditions were recorded on samples containing 300 μM insulin in the absence and presence of 1x molar ratio GM3 in a buffer containing 10 mM NaPi pH 3.0, 100 mM NaCl, and 10% D_2_O. Spectra were acquired with 32,768 points, 256 scans, 16.23 ppm spectral width, 3290.00 Hz transmitter frequency offset, and a recycle delay of 1.0 s. All NMR spectra were processed with 8,192 effective points (TD_eff_) and a size of the real spectrum of 131,072 points. A gaussian window function was applied (LB: -5.0 Hz, GB: 0.05) and the baseline was corrected using *qfil* with a filter width of 0.4 ppm.

### 2.8. Filtration of Insulin-Ganglioside Samples

Centrifugal ultrafiltration of insulin reaction samples was performed using Amicon Ultra 2 mL centrifugal filters with a 3 kDa molecular weight cutoff (MilliporeSigma). Prior to sample loading, the filter membranes were pre-rinsed with 600 μL of 10 mM sodium phosphate buffer at pH 3 by centrifugation at 4500g for 20 minutes. The insulin concentration in the insulin-ganglioside samples was measured using a NanoPhotoMeter NP80 (IMPLEN). After rinsing, 240 μL of each sample was added to the upper reservoir of the filtration units and centrifuged at 4500 g for 10 minutes at 4 °C. Once the volume of the sample was reduced to approximately 50 μL, the retentate was diluted back to the original volume with fresh buffer and centrifuged again. This washing step was repeated 3-4 times for each sample to ensure effective removal of the unbound gangliosides. After the final spin, the retentate was collected in clean tubes, diluted, and the final insulin concentration was remeasured using the NanoPhotoMeter NP80 (IMPLEN). Sialic acid assay to quantify the amount of unbound ganglioside being filtered out was performed with Sialic Acid Assay Kit (Sigma-Aldritch, Catalog # MAK314-1KT), according to the manufacturer’s instructions.

### 2.9. Cell toxicity assays

Cellular toxicity in NIH3T3 mouse fibroblast cells was assessed using the XTT cell viability assay. NIH3T3 cells were cultured in Dulbecco’s Modified Eagle medium (DMEM) supplemented with 10% fetal bovine serum (FBS) and 1% penicillin-streptomycin, maintained in a humidified incubator at 37 °C with 5% CO_2_. Cells were subcultured upon reaching 70%-80% confluency. The culture medium was removed using glass pipettes and vacuum, and the cells were washed with phosphate-buffer saline (PBS). A 0.25% trypsin-EDTA solution was added to detach the cells, followed by incubation for 5-6 minutes. The detached cells were centrifuged, resuspended in fresh culture medium, and transferred to new culture vessels to ensure optimal growth conditions. Cells were subcultured a minimum of three splitting cycles after revival from frozen stock prior to the toxicity experiments. The XTT assay was performed using the CyQUANT™ XTT Cell Viability Assay (Thermo Fisher Scientific, Catalog # X12223). A total of 15000 cells were added to wells in a transparent 96-well plate in 100 μL of cell growth medium. After 24 hours of incubation, the cells were treated with insulin reaction samples and allowed to incubate for an additional 24 hours. Per the manufacturer’s protocol, 1 mL of Electron Coupling Reagent was added to 6 mL of XTT Reagent and mixed well. 70 μL of this working solution was added to each well. Then the plate was incubated for 4 hours at 37 °C and the absorbance was measured at 450 nm and 660 nm to evaluate cell viability. Cell viability data were analyzed using one-way analysis of variance (ANOVA) followed by Tukey’s Honestly Significant Difference (HSD) post hoc test to determine statistically significant differences between insulin-treated samples and ganglioside co-treatment groups. Three biological replicates were used for each condition. The analysis was performed in MATLAB using built-in functions (anova1) and (multcompare). For each condition, only pairwise comparisons with the insulin-alone treatment group were extracted and reported. A significance threshold of *p* < 0.05 was used. Significance levels were denoted using asterisks as follows: *p* < 0.05 (\**), p < 0.01 (**), and p < 0.001 (\*\*\**). Comparisons with *p* ≥ 0.05 were considered not significant (ns).

### 2.10. Seeding Activity of Insulin-Ganglioside Aggregates

Insulin-Ganglioside seeds were prepared by centrifuging 200 μL of the insulin+5× ganglioside samples at 14000 rpm for 15 minutes at 4□°C in a Sorvall ST 16R (ThermoFisher Scientific). After centrifugation, the supernatant was removed and replaced with 200 μL of deionized Milli-Q water. This wash step was repeated three times to remove the residual unbound components. The isolated seeds were then added to freshly prepared 80 μM insulin solution with 150 mM NaCl in 12.5%, 25%, 37.5% concentrations and then incubated for 48 hours.

### 2.11. Small Angle X-ray Scattering

SAXS experiments were performed on a Xenocs Xeuss 3.0 SAXS instrument. The X-ray wavelength was 1.34 Å; and the sample-to-detector distance was 900 mm. The X-ray beam size was 0.5 mm in diameter. Liquid samples were loaded in quartz capillaries with a diameter of 2 mm, sealed with wax, and measured in vacuum environment. X-ray exposure time was 20 min. 1D data fitting was carried out using Irena macro based on Igor software [52].

### 2.12. Molecular Dynamics Simulation

All molecular dynamics simulations were performed using version 8.1.2 of the OpenMM simulation platform using the Charmm36 forcefield [53,54]. The three-dimensional structure of human insulin was obtained from the Protein Data Bank with ID 1EV3 [55]. From the structure, only the monomeric form consisting of the A and B chains was used in simulations. For simulations performed at pH 3, insulin was protonated at the residues Hsp-5, Hsp-10, Glu-13, and Glu-21 of the B chain. GM3 and GD3 structures were constructed with the aid of Charmm-GUI, which was used to assemble lipid bilayers containing 80 lipid molecules in each leaflet of the bilayer [56,57]. An additional 27761 water molecules using the TIP3P forcefield were added in the simulation resulting in average simulation dimensions of 7.36×7.30×20.5 nm for GM3 and 8.25×8.26×16.5 nm for GD3. Sodium and chlorine ions were added as necessary to maintain electroneutrality while achieving an ionic strength of 150 millimolar in accordance with experiment.

Insulin and the bilayers were initialized under the standard Charmm-GUI initialization and equilibration where the solvent, then lipids, then protein sidechains, then protein backbone sequentially have their constraints removed and are individually allowed to equilibrate. A small external potential of the form 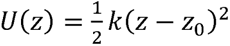 with *k* equal to 0.1kJ/mole-nm^2^ and *z_0_* equal to 9.5 nm was applied only to the lipid molecules in order to bring both leaflets together before being turned off for full equilibration and production. All simulations were performed using the Langevin integrator with a time step of 0.002 ps, with a frequency of 1 ps^-1^ at a temperature of 310 K. In order to allow the membrane to equilibrate, a Monte Carlo membrane barostat was used with a pressure of 1 bar and a membrane surface pressure of 0 bar-nm. All systems were after energy minimization allowed to freely relax for 75 ns before starting umbrella sampling [58]. For umbrella sampling a series of eight simulations were conducted for each lipid and pH combination to determine the potential of mean force with center of mass distance along the *z* coordinate using the potential

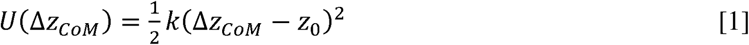

where Δz_CoM_ is the distance along the z coordinate between the protein and bilayer center of mass. The spring constant k was set to 10 kJ/mole-nm^2^ while z was varied between 3.5 nm and 6.0 nm via eight evenly spaced points. Umbrella sampling simulations were allowed to equilibrate for 50 ns before 100 ns of production.

## 3. Results

### 3.1. Gangliosides Accelerate Aggregation of Insulin

To investigate the impact of ganglioside lipids on the aggregation kinetics of insulin, ThT fluorescence assays were performed using 80 μM insulin at 37° C under continuous shaking at pH 3 without gangliosides or in the presence of GD3/GM3 ganglioside with concentrations ranging from equimolar to fivefold molar excess relative to the insulin concentration (Fig. 1). The ThT fluorescence data revealed that the presence of both GM3 and GD3 augment insulin aggregation significantly with increasing molar ratio of lipids as compared to insulin aggregation without any lipids (s. 1). Additionally, sigmoidal fitting of the kinetics curves revealed that ganglioside lipids significantly reduced the lag time (t_lag_) and half-time of aggregation (t□/□) of insulin aggregation in a concentration dependent manner (Table. S1). In contrast, in absence of lipids, insulin exhibited a longer lag time of approximately 34 hours. At equimolar concentrations, the lag times were reduced to approximately 18 hours for GD3 and 25 hours for GM3. With increasing concentration of gangliosides, the lag time decreased further (Fig.1, Table S1). The shortest lag times were observed at the highest concentrations of gangliosides, around 9 h for GD3 and 10 h for GM3. It is worth noting that although the shortest lag time is similar for both GD3 and GM3, for the 1.5× and 2.5× concentrations of these lipids, the lag times do vary between the two lipids. Evidently, GD3 exhibited consistently shorter lag times compared to GM3 at corresponding concentrations. Overall, these results demonstrate that both GD3 and GM3 modulate insulin aggregation kinetics in a concentration-dependent manner, as evidenced by their ability to progressively shorten the lag phase of aggregation.

**Figure 1.**
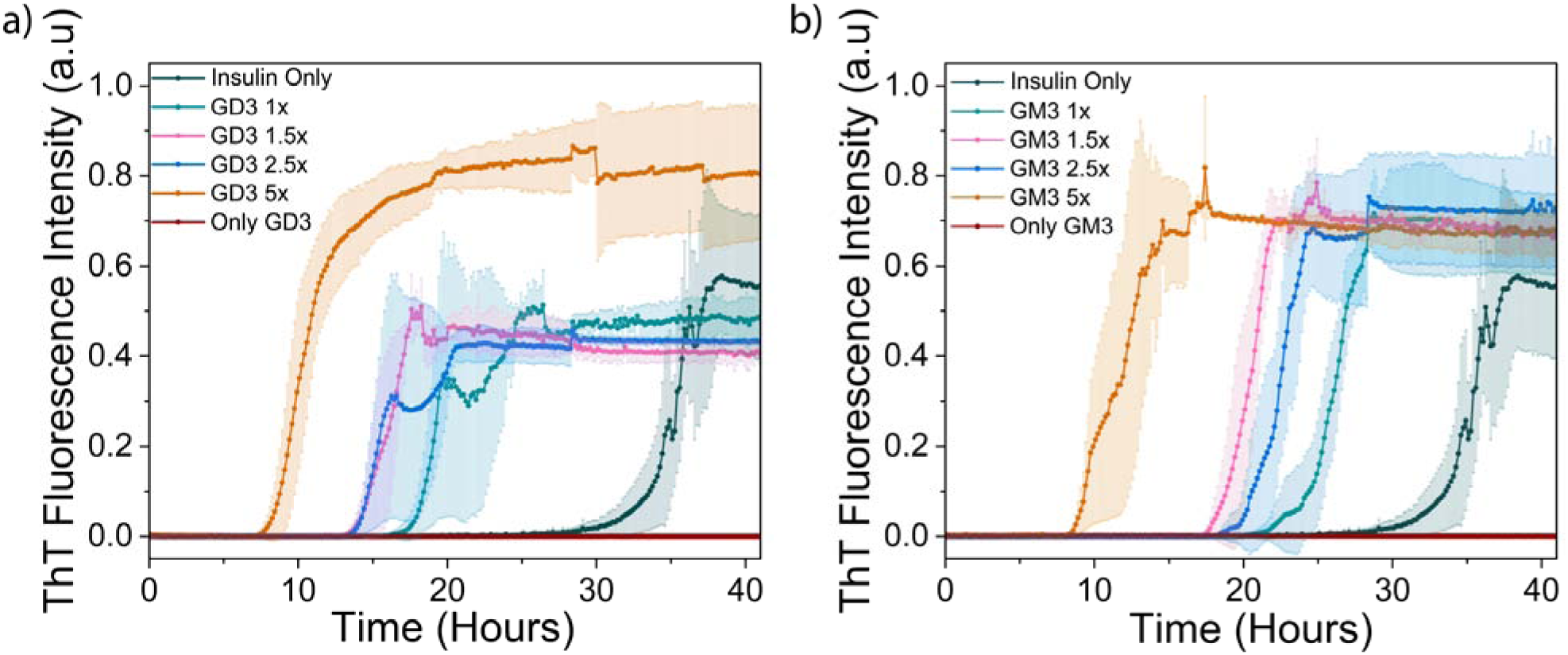
Thioflavin T (ThT) fluorescence kinetics of 80 μM insulin in the absence (dark green) or presence of gangliosides at increasing concentrations relative to insulin: 1× (green), 1.5× (pink), 2.5× (blue), and 5× (yellow). Data are shown for (a) GD3 and (b) GM3. Reactions were carried out in 10 mM sodium phosphate buffer (pH 3.0) containing 150 mM NaCl and 10 μM ThT, at 37 °C under continuous shaking at 700 rpm. ThT fluorescence from lipids alone samples are shown in red. Insulin monomers were freshly prepared in the same buffer immediately prior to initiating the reactions.

### 3.2. The presence of gangliosides alters the secondary structure and morphology of insulin aggregates

To investigate the effects of gangliosides GM3 and GD3 on insulin’s morphology and secondary structure, a combination of Fourier-transform infrared (FTIR) spectroscopy, circular dichroism (CD) spectroscopy, small angle x-ray scattering (SAXS) and transmission electron microscopy (TEM) was employed. TEM micrographs of insulin aggregates at 42 h showed that insulin fibrils formed in the absence of gangliosides were homogeneous with long, unbranched structures exceeding 100 nm in length (Fig. 2). In contrast, the presence of GM3 or GD3 resulted in a pronounced effect in aggregate morphology of insulin. At all tested concentrations, the characteristic fibrillar structures were absent. Instead, the aggregates displayed a uniform bead-like morphology which resembled discrete globular subunits (Fig. 2). These results imply that gangliosides interfere with the canonical fibrillation pathway of insulin, promoting the formation of structurally distinct, non-fibrillar aggregates. Similarly, a lack of mature fibril morphology was also observed with insulin aggregates formed in the presence of GM3; however, subtle distinctions in insulin morphology were observed at lower GM3 concentrations as compared to insulin aggregates formed in the presence of GD3. Specifically, at 1× and 1.5× GM3, the aggregates included short and curved, fibril-like morphology indicating the presence of pre-fibrillar intermediates. However, these structures lacked the length, and uniformity of insulin amyloid fibrils formed in the absence of gangliosides. These results indicate partial progression along the aggregation pathway before redirecting towards less-ordered, amorphous assemblies.

**Figure 2.**
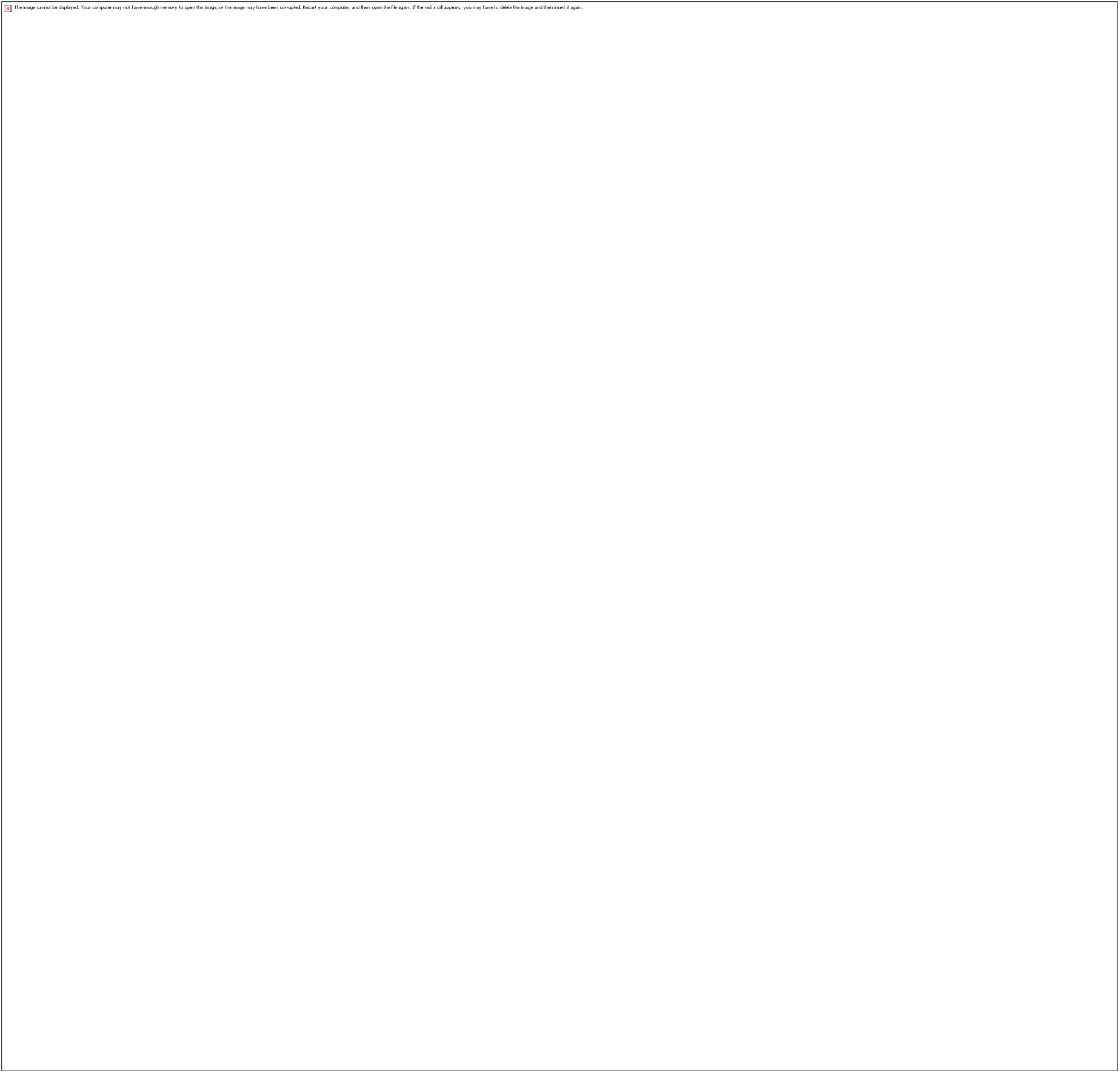
TEM images showing the effect of gangliosides GD3 and GM3 on insulin aggregate morphology. yield short, beaded structures whereas ganglioside-free insulin forms fibrils. TEM Images of (a) Insulin forms long, unbranched fibrils in the absence of lipids. Insulin incubated with GD3 at concentrations of 1× (b), 1.5× (c), 2.5× (d), and 5× (e) exhibit short, beaded, non-fibrillar structures. Corresponding TEM images for insulin with GM3 1× (f), 1.5× (g), 2.5× (h), and 5× (i) show similar beaded morphologies. Indicated scale bars are 90 nm(a), 100 nm (b-d, f-h) and 200 nm (e,i). Small-angle X-ray scattering (SAXS) analysis of insulin aggregates formed with gangliosides GD3 and GM3(j-m). SAXS profiles of insulin aggregates formed in the presence of GD3 or GM3 at 1× (j) and 5× (k) molar ratios compared to insulin alone. Trial fittings show that at 5× concentrations, GD3 and GM3 samples exhibit distinct Bragg peaks(l,m), indicating the formation of internally ordered, non-fibrillar assemblies, in contrast to the fibrillar aggregates observed in lipid-free conditions.

Small-angle X-ray scattering (SAXS) measurements were conducted to investigate the nanoscale packing of insulin aggregates formed in the presence and absence of gangliosides. SAXS curves are shown in Fig. 2, which helps identify large-scale aggregated structures as well as local packing of insulin-ganglioside aggregates. All curves show an intensity upturn, following a power law with scaling exponent between 3 and 4 in the low-q region (q < ∼0.005 Å□¹); such features are due to large-scale structural inhomogeneities, extended beyond the probing range of SAXS, > ∼100 nm. In the intermediate-q range between ∼0.006 and 0.2 Å□¹, distinct local structures in the order of 10 nm were observed. Intriguingly, at higher concentrations of lipid, namely, for GM3 5× and GD3 5× samples, prominent Bragg’s peaks can be identified, related to packing of gangliosides into macro-lattice. The position of the first peak of GM3 5× and GD3 5× samples are 0.071 and 0.085 Å□¹, corresponding to d-spacings of 8.3 and 7.3 nm, respectively. Trial fittings indicate that body-centered cubic (BCC) and double diamond (DD) cubic phase can be used to index all Bragg’s peaks (see the solid lines and indicated peak positions in Fig. 2 (l,m). For GM3 5×, the peak positions follow a ratio of 1:√2:√3… (BCC). Peaks in GD3 5× appear at larger angles, and their separations are clearly lesser, which seems to follow a ratio of √2:√3:2…(DD). Due to structural disorder and size effect, those peaks are severely broadened, precise determination of the space group becomes challenging, which may be resolved in future studies when more perfect macro-lattices can be prepared for structure determination purposes. The dimensions for the two cubic cells of GM3 5× and GD3 5× are 12.5 and 10.4 nm, respectively. These findings imply that at higher lipid concentrations, gangliosides promote the formation of structured, non-fibrillar amorphous assemblies with periodic internal packing, in contrast to the fibrillar aggregates observed in lipid-free conditions. Thus, the findings from both TEM and SAXS highlight that GD3 and GM3 distinctly influence insulin aggregation, leading to subtle variations in the aggregation pathways and resulting in structurally different aggregate morphologies.

FTIR spectra of insulin fibril aggregated without gangliosides displayed a prominent peak at ∼1633 cm□¹, indicative of β-sheet structure [59], along with a peak near 1667 cm□¹ corresponding to β-turns [60] (Fig. 3). In contrast, a β-sheet peak accompanied by a shoulder near 1655 cm□¹ was observed in insulin aggregates formed in the presence of gangliosides, indicating an increase in α-helical content. To identify and quantify the structural components of the insulin aggregates, peak deconvolution of FTIR spectra was performed using Gaussian equation in Origin (Table S2). The insulin fibrils generated without gangliosides exhibited 47.6% β-sheet and 48% β-turn content, with minimal (3.83%) α-helix content, consistent with the TEM data of mature amyloid fibrils. Additionally, aggregates formed in the presence of the lowest concentration of GM3 did not exhibit any helical structure. However, in the presence of 5× GD3 the aggregates displayed 73.9% β-sheet structure, reduced β-turns, and a notable 11.8% α-helical content. Similarly, 1.5× GD3 resulted in a remarkably high β-sheet content (87.5%) and minimal β-turns with a 3.6% α-helical content. In contrast, insulin aggregation with 1× or 2.5× molar concentrations of GD3 led to a significant increase in α-helical structure (∼ 50%) accompanied by a decrease in β-sheet and β-turn structures. Conversely, the presence of GM3 resulted in a concentration-dependent increase in α-helical content, with 41.6% α-helix in the presence of 5× GM3. To further validate these findings, the CD spectra were acquired for the same samples (Fig. 3, Table S3). At 42 hours, the insulin fibril sample generated without gangliosides displayed a single minimum at around 218 nm, characteristic of β-sheet-rich structures. In contrast, insulin samples aggregated with gangliosides displayed a double minimum pattern with two minima at 208 and 222 nm, suggesting an increase in α-helical content. At 9-hour, CD spectra of all samples show α helical structure, but at 16-hours, the insulin only sample can be seen starting to form β-sheet structures (Fig. S2). Further deconvolution of CD spectra was consistent with this interpretation of the absence of α-helical content in the insulin control and an increased α-helical content in the insulin samples with gangliosides (Table S3). The highest α-helical content (7.4%) was present in the sample with 5× GM3, corroborating the FTIR results. Taken together, the results from FTIR and CD indicate that gangliosides, particularly GM3, modify the secondary structure landscape and retain a substantial proportion of α-helical content of insulin, particularly at higher concentrations of ganglioside. This, in contrast with the insulin fibrils generated without gangliosides which features mostly β-sheet structures, suggests that gangliosides, especially GM3, may stabilize native-like or intermediate helical conformations during aggregation.

**Figure 3.**
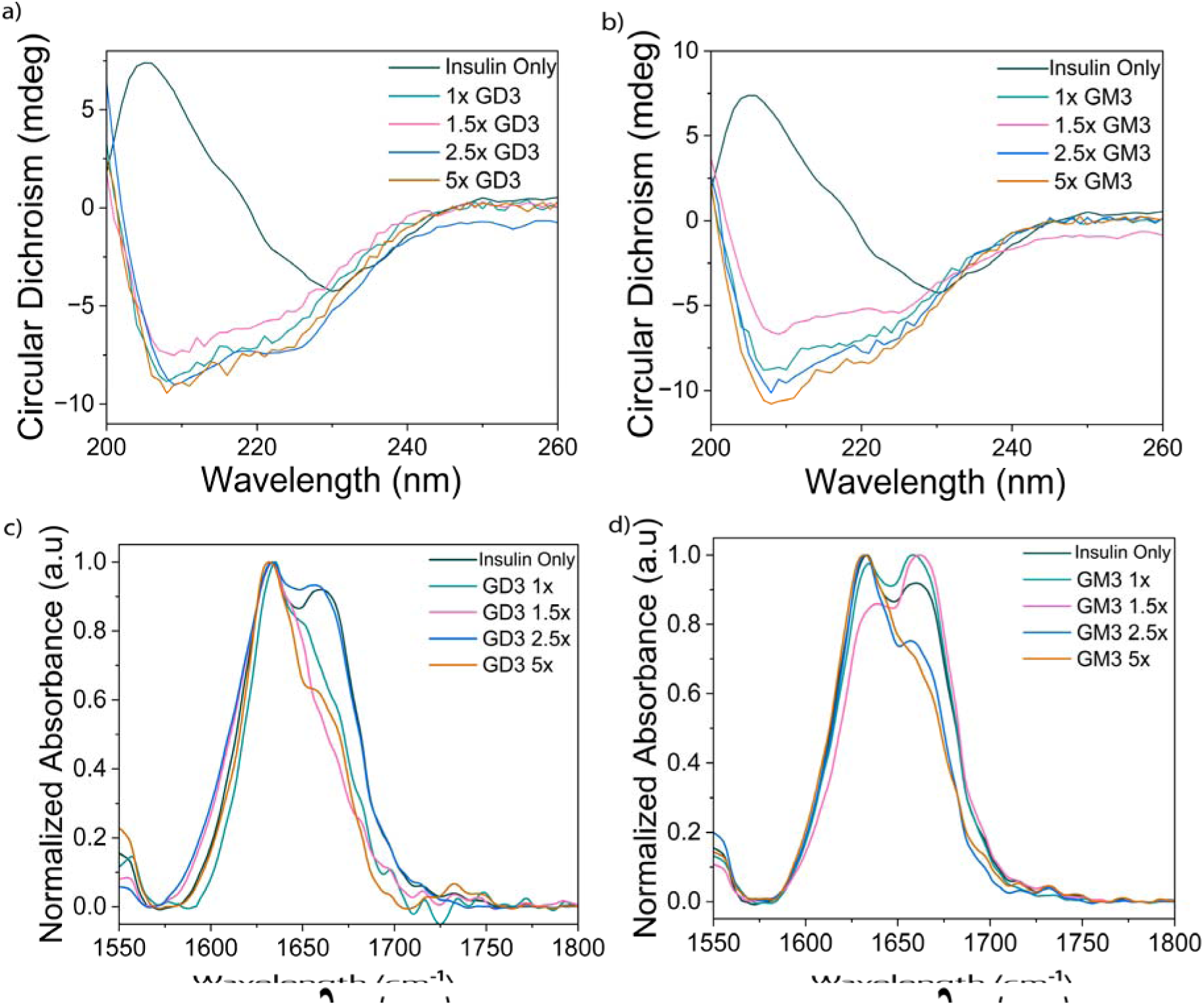
(a,b) CD spectra and (c,d) FTIR spectra of 80 μM insulin in a lipid free environment (dark green) and in the presence of increasing concentrations of gangliosides, GD3 (a and D) and GM3 (b and c): 1× (green), 1.5× (pink), 2.5× (blue), and 5× (yellow). (a) GD3 and (b) GM3 after 42 hours of incubation.

### 3.3. Interaction between gangliosides and insulin

To better understand the interaction of the insulin monomer with GM3 and GD3, molecular dynamics simulations were performed. In these simulations, GM3 and GD3 were treated as forming planar bilayers which interact with the insulin molecule. Systems simulated were insulin at pH=3.0 and pH=7.0 in contact with either a pure GM3 or pure GD3 bilayer (Fig. S3). Upon equilibration it was noted that the GD3 lipids had a larger area per lipid of 0.849 nm^2^/molecule compared to GM3’s 0.678 nm^2^/molecule. Two different conformations of the insulin monomer were simulated: the first the native monomer conformation, the second a denatured conformation obtained by applying an external harmonic potential to the B chain a-helix to cause it to unfold. It was noted that upon unfolding a small b-sheet region formed between the A and B chains as shown in Fig. 4.

**Figure 4.**
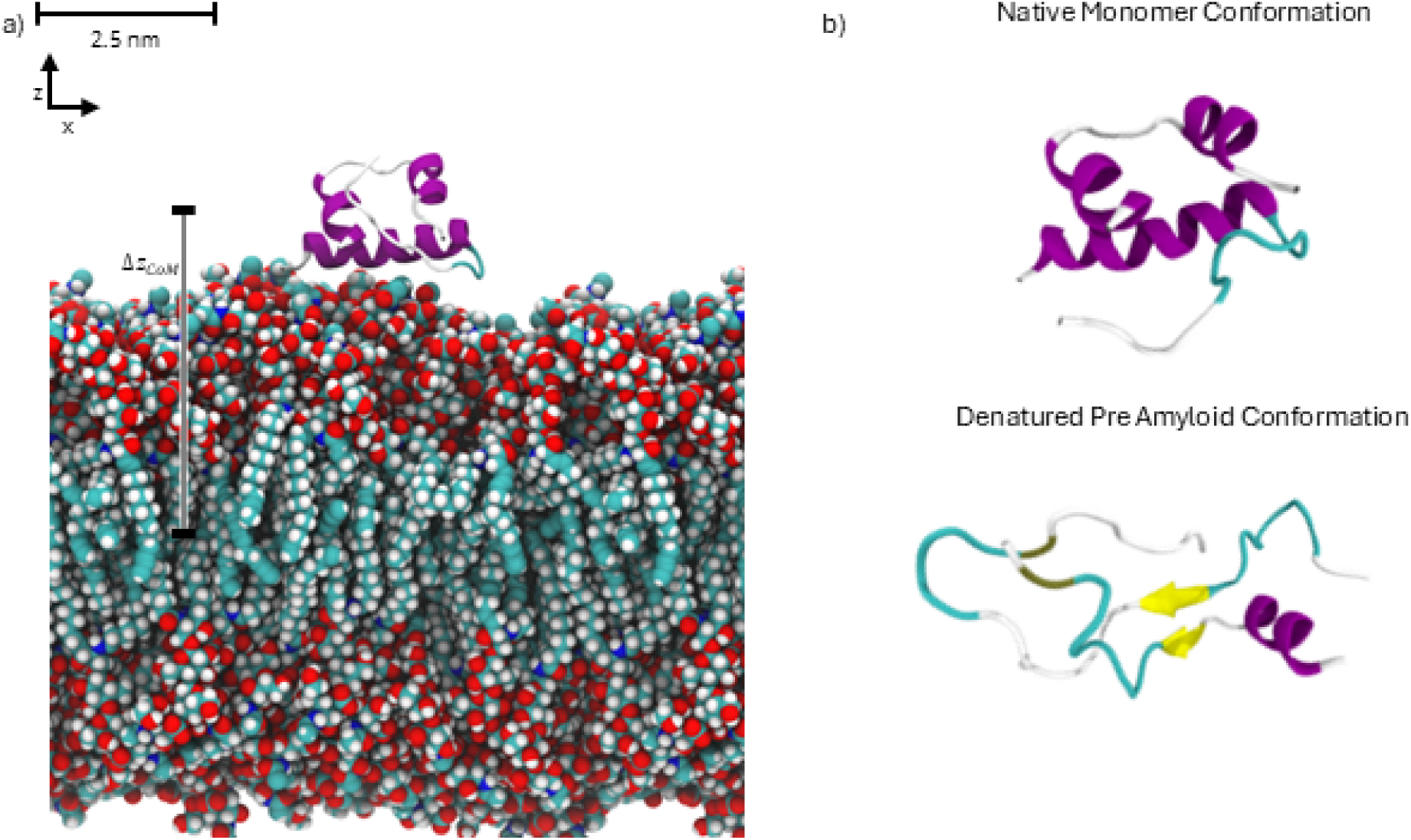
(a) Simulation scheme for an insulin molecule in contact with GM3 lipid bilayer. Center of mass distance along the z coordinate measured between the center of the bilayer and the center of mass of the insulin molecule. Simulation scheme for an insulin molecule interacting with GD3 bilayer is shown in Fig. S4. (b) Representative conformations for the folded and unfolded insulin molecules used in th simulation studies. Color code: Magenta – Alpha Helix, Cyan – Turn, Yellow – Beta Sheet, Green – Sulfide bridge.

In order to examine the free energy of insulin associating with the lipid bilayer’s surface, a series of umbrella sampling simulations was carried out in which the insulin molecule was placed in a harmonic potential with varying equilibrium distances from the lipid bilayer center z coordinate as depicted in Fig. 5. Resulting probability distributions of the distance were used to construct the free energy surface using the multistate Bennett acceptance ratio (MBAR) method [61,62]. Results for the potential of mean force are shown in Fig. 5. In all cases for the folded conformation, there exists a minimum in the potential of mean force at a z coordinate center of mass separation between 4.0 nm and 4.5 nm, which corresponds to the protein absorbed on the bilayer surface. The depth of this well is estimated to be between 2 to 4 kJ/mol corresponding to a moderate partitioning of the insulin molecule to the lipid bilayer surface from solution while still able to freely desorb. Across simulations performed, GM3 and GD3 exhibited similarly shaped potentials of mean force. We found that protonation of the insulin molecule at pH 3.0 leads to increased absorption, which is hypothesized to be due to increased polar interactions of the insulin molecule with the interface. This is further substantiated by the salt-dependent aggregation behavior of insulin observed in NMR experiments (*vide infra*, Fig. 7). In the absence of salt, insulin readily forms visible aggregates with increasing concentration of ganglioside, while this effect is reduced in the presence of 100 mM NaCl. In the case of the unfolded insulin, we find that the depth of the well is much less pronounced than in the folded case and is spread over a larger distance from the interface. We hypothesize that this change in energetics is due to increased flexibility of the insulin molecule in the denatured state, allowing for polar residues to remain in contact with the interface over larger distances than in the folded conformation.

**Figure 5.**
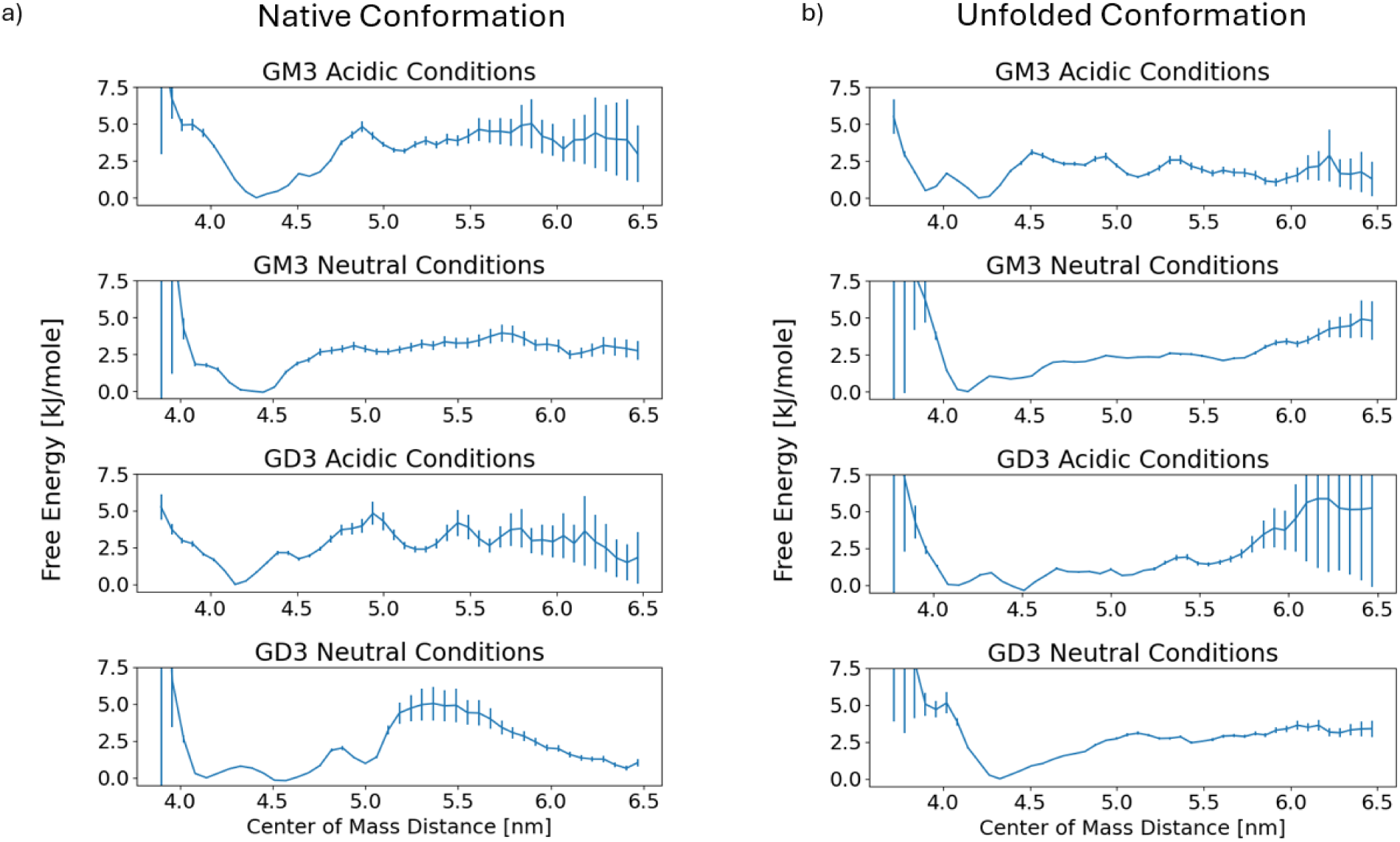
(a) Potential of mean force of the native conformation protein/bilayer system as a function of z coordinate distance between the bilayer center of insulin center of mass. (b) Potential of mean force of the unfolded protein/bilayer system as a function of z coordinate distance between the bilayer center of insulin center of mass.

To further examine the binding behavior, the contact frequency was calculated between insulin residues and the lipid surface, where a residue in contact is defined as any residue with its atom closer than 2.5 angstroms from an atom in a lipid molecule. We find that beyond a z distance of 5.5 nm; the insulin molecule has no residues in contact with the interface in the folded case. In contrast the unfolded molecule may remain in contact with the interface up to 5.5 nm of separation. Results for the overall contact frequency of the residues with the bilayer are shown in Fig. 6. Under acidic conditions, for the folded protein, the residues along the B chain alpha helix are the primary contacts with the lipid bilayer, including the residues Phe-1-B, Val-2-B, Asn-3-B, Hsp-5-B, and Hsp-10-B. Notably the residues Hsp-5-B and Hsp-10-B are protonated under acidic conditions leading to increased interactions with the negatively charged lipid bilayer. Under neutral conditions residues in contact with the lipid bilayer are less specific, however Phe-1-B remains in contact, with the other most frequent contacts being Phe-1-B, Leu-16-A, Gly-1-A, and Typ-14-A. Similar trends are observed for the unfolded state. However, due to the absence of the alpha helix and more flexible conformation, contacts are less specific.

**Figure 6.**
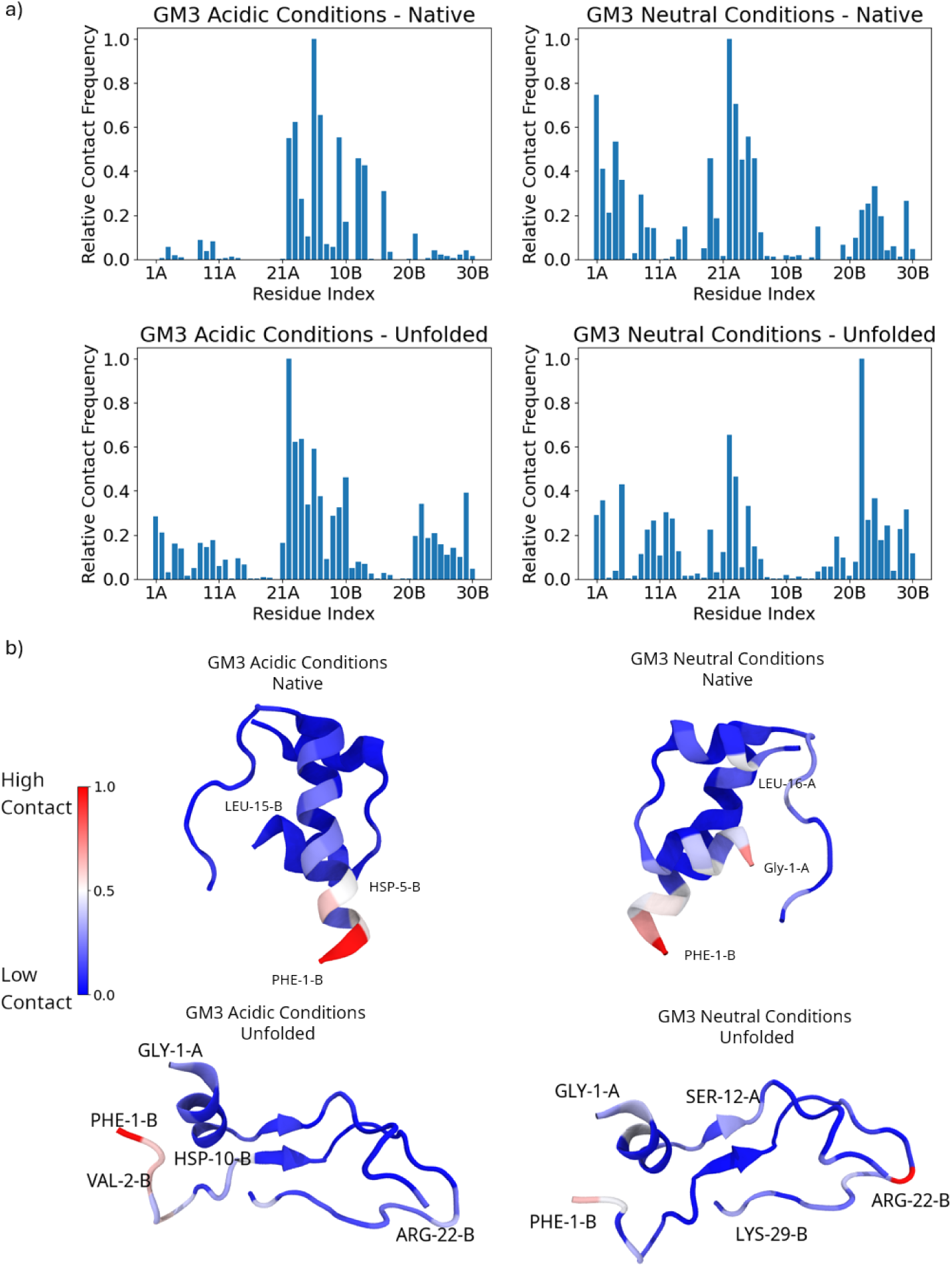
(a) Plots of contact frequency between each residue of the insulin and lipid bilayer normalized to the most frequent in contact residue. (b) Insulin monomers with residues are colored by overall contact frequency: red as the most frequent and blue as the least frequent. Data for GD3 are shown in supplementary information (Fig. S5). Videos demonstrating representative molecular dynamics trajectories illustrating insulin rotational dynamics when bound to ganglioside-containing lipid bilayers under different lipid compositions and pH conditions are shown in SV1.

This lack of specificity under neutral conditions is hypothesized to be due to a lack of positively charged residues under neutral conditions that promote interaction with the negatively charged lipid bilayer. We note that because the umbrella sampling simulations were only allowed to run for 100 ns of production and initialized from the properly folded insulin structure, they are only sufficient to describe the interaction of the folded insulin protein with the polar outer surface of the lipid bilayer. The umbrella sampling simulations further were only biased in the protein-bilayer distance, and so did not explicitly target changes in conformation that could occur while in contact with the interface.

In addition to the molecular simulations, solution NMR experiments were performed to probe the molecular interactions between gangliosides and insulin (Fig. 7). Exposing insulin to 1x GM3 under low-salt conditions resulted in immediate, visible precipitation which is also evidenced by a loss in signal intensity of approximately 30% in 1D ^1^H NMR spectra (Fig. 7A). Under these conditions, increasing the lipid concentration to 5x GM3 resulted in further, heavy precipitation and the loss of all insulin signals. Notably, when insulin was exposed to GM3 in a 1:1 ratio under salty conditions (100 mM NaCl, Fig. 7B), less precipitation was observed and the loss in signal intensity was reduced to approximately 17%. These results point to a charged interaction between the positively charged insulin and the negatively charged GM3 micelles, where insulin molecules are readily absorbed by the surface of the GM3 micelle. During this process, large aggregates are formed that are visible to the naked eye and rapidly sediment in the NMR tube. The NMR spectra show no evidence of the presence of GM3, neither in the form of broad signals stemming from micellar GM3 nor sharp signals resulting from free monomeric GM3. This is in line with previous studies indicating the formation of large micelles where GM3 signals are broadened beyond detection [63]. An alternative explanation for the absence of GM3 signals in 1D NMR spectra is that GM3 and insulin form a co-precipitate where the majority of GM3 is removed from solution. Compared to GM3, when insulin is exposed to GD3 in a 1:1 ratio under identical, salty conditions (100 mM NaCl, Fig. 7C), significant precipitation was observed and approximately 80% of signal intensity was lost compared to a loss in 17% for GM3. This is in line with observations in the ThT assays where it has been exhibited that GD3 causes faster aggregation in insulin compared to GM3. These observations underline the role of charged interaction between the positively charged insulin and the negatively charged ganglioside micelles. The rapid aggregation observed in NMR spectroscopy appears to be directly correlated to the number of negative charges on the surface of the ganglioside micelle, with one negative charge per lipid molecule in GM3 micelles (mono-sialic acid) and two negative charges per lipid molecule in GD3 micelles (di-sialic acid).

**Figure 7.**
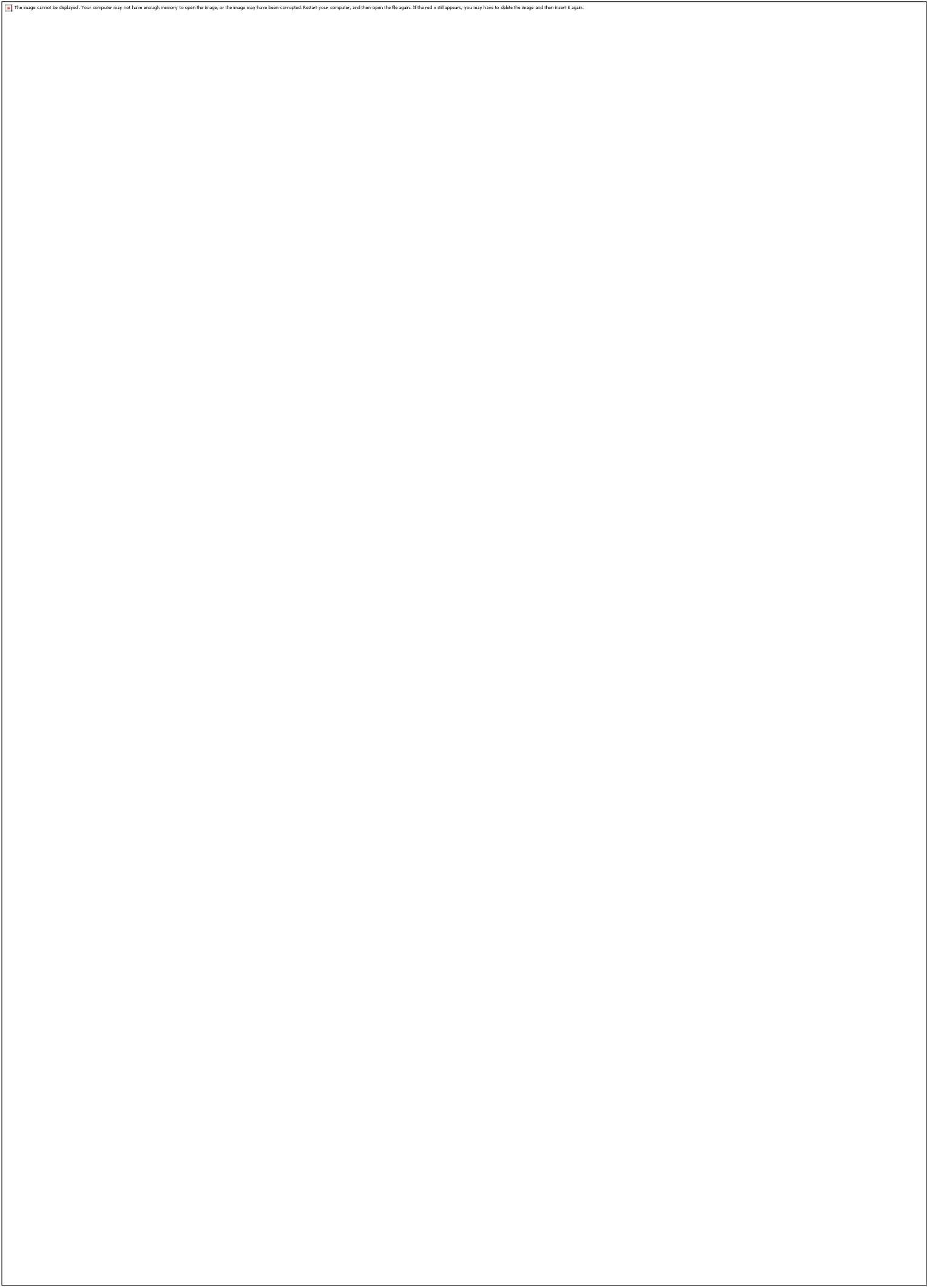
Interaction of insulin with GM3 and GD3 using NMR spectroscopy. (a) ^1^H NMR spectra of a sample containing 80 μM insulin under low-salt conditions (10 mM NaPi pH 3.0, 10% D_2_O) in the absence (dark green) and presence of 1x GM3 (light green) and 5x GM3 (orange). Residual peaks in the orange spectrum belong to small-molecule impurities. (b) ^1^H NMR spectra of a sample containing 300 μM insulin under salty conditions (10 mM NaPi pH 3.0, 100 mM NaCl, 10% D_2_O) in the absence (dark green) and presence of 1x GM3 (light green). (c) ^1^H NMR spectra of a sample containing 300 μM insulin under salty conditions (10 mM NaPi pH 3.0, 100 mM NaCl, 10% D_2_O) in the absence (dark green) and presence of 1x GD3 (light green). Truncated peaks in the light green spectrum belong to small-molecule impurities. All spectra were recorded at 700 MHz and 298 K.

### 3.4. Gangliosides mitigate the cytotoxicity of insulin aggregates

To assess whether changes in aggregate structure induced by gangliosides affect cellular toxicity, insulin-ganglioside species were incubated with NIH3T3 mouse fibroblast cells, and XTT cell viability assay was performed (Figs. 8, S6). For initial assessment of cytotoxicity, the insulin-ganglioside aggregates samples (with 80 μM insulin) generated with 5× or 1× molar excess of gangliosides, which included both the aggregates and any unbound gangliosides. The XTT assays revealed that the insulin fibrils were highly toxic, for both 40 μM and 5 μM doses (insulin concentration), compared to insulin-ganglioside aggregates confirming the toxicity of insulin fibrillar species (Fig. 8). However, aggregates formed in the presence of GM3 and GD3 substantially reduced cell toxicity at 5 μM, with cell viability comparable to the untreated controls. But this protective effect was not seen in 40 μM doses. To understand whether this reduction in toxicity was due to structural changes in the aggregates or the presence of unbound gangliosides in the samples, we analyzed the cellular toxicity of filtered samples. The insulin-ganglioside aggregates were filtered using Amicon Ultra-2 mL centrifugal filters with a molecular weight cutoff of 3 kDa, which removed unbound gangliosides in the samples (Fig. S6) (verified by sialic acid assay). Interestingly, the filtered samples of insulin-ganglioside aggregates showed improved cell viability even at 40 μM. This indicates that the presence of free gangliosides did not mitigate cell toxicity and may even have contributed to the reduced cell viability in the high concentration doses. The XTT assay of the ganglioside only controls on NIH3T3 cells revealed that the gangliosides reduced cellular toxicity indicating that exogenous addition of gangliosides are not inherently protective in nature and may exert toxicity at higher concentrations (Fig. S6) as reported previously [64–67]. Taken together, these findings suggest that GM3 and GD3 reduce the toxicity of insulin aggregates primarily by altering their secondary structure and morphology, rather than by directly protecting cells as free lipids.

**Figure 8.**
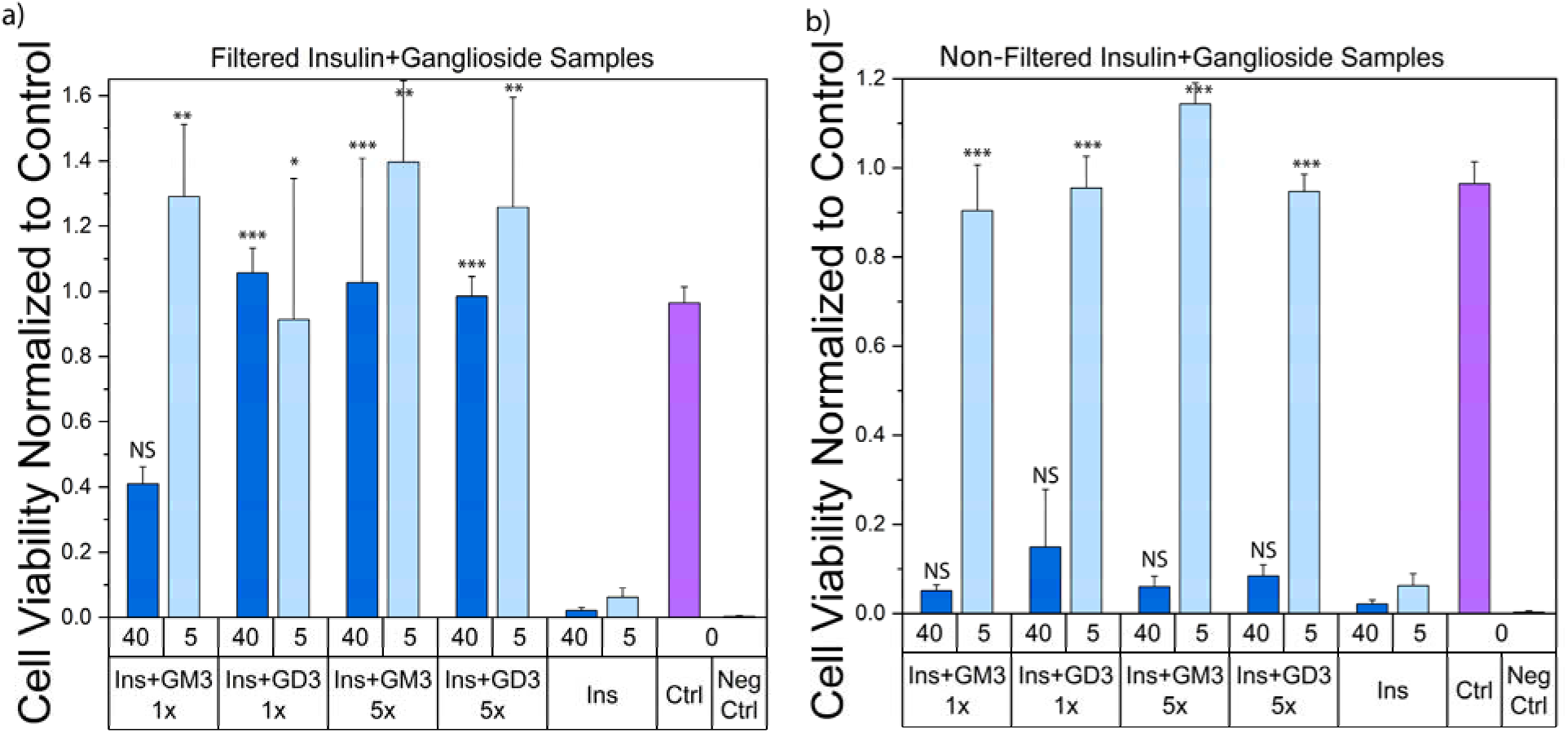
Viability of NIH3T3 cells incubated with (a) non-filtered and (b) filtered insulin-ganglioside aggregates. Cell viability was assessed using the XTT cell viability assay for NIH3T3 cells after 24 hours of incubation with the indicated samples. Non-filtered samples contained both aggregates and unbound gangliosides, while filtered samples contained only aggregates. Statistical significance was determined by one-way ANOVA followed by Tukey’s post hoc test, comparing each treatment group to the insulin-alone condition. Asterisks denote significance levels: *p* < 0.05 (\**), p < 0.01 (**), and p < 0.001(\*\*\**), and ns = not significant (*p* ≥ 0.05).

### 3.5. Gangliosides do not influence the aggregation of reduced insulin

To assess how gangliosides GD3 and GM3 affect the aggregation of reduced insulin, ThT fluorescence over 48 hours in the presence and absence of these lipids were monitored (Fig. 9). Reduced insulin alone exhibited minimal ThT fluorescence, indicating a lack of classical amyloid formation. Interestingly, the addition of GD3 and GM3 did not alter this profile. Instead, the ThT signal remained low and flat, compared to the insulin control, closely matching that of the reduced insulin control. This observation suggests that both GD3 and GM3 gangliosides do not promote or restore ThT binding structures under reduced conditions. Consistent with the ThT fluorescence results, TEM micrographs of these species revealed amorphous aggregates with no discernible fibrillar morphology (Fig. 9). These results suggest that, unlike in non-reducing conditions, gangliosides GD3 and GM3 do not affect the aggregation of reduced insulin and do not induce a transition toward fibrillar structures, rather retain its non-fibrillar aggregation pathway.

**Figure 9.**
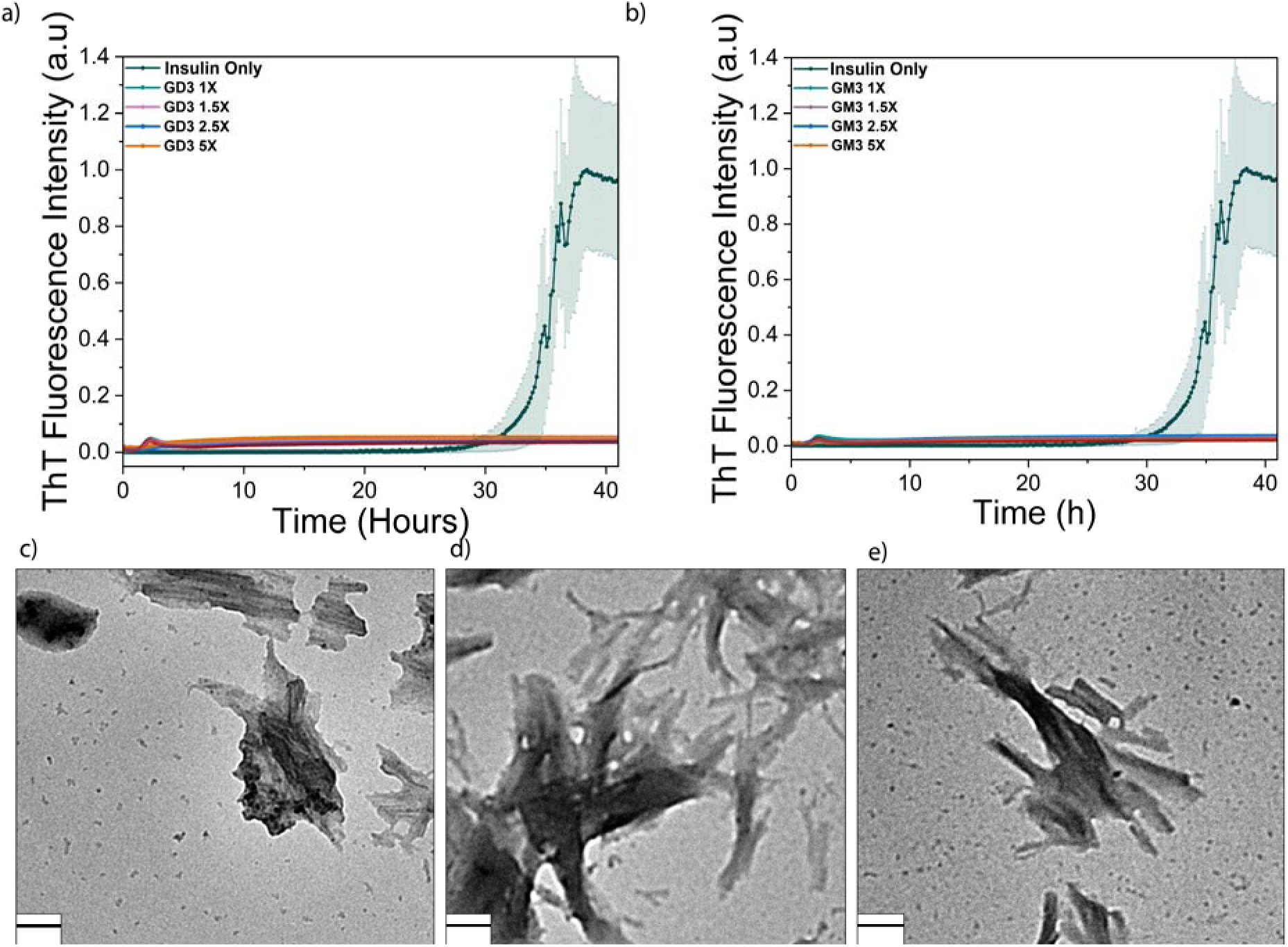
ThT fluorescence kinetics of 80 μM insulin reduced by TCEP HCl in a lipid free environment (yellow), and in the presence of increasing concentrations of gangliosides GD3 (a) and GM3 (b): 1× (green), 1.5× (pink), 2.5× (blue), and 5× (yellow). Non-reduced insulin monomers (dark green) are shown for comparison. Insulin and gangliosides were incubated in 10 mM sodium phosphate buffer (pH 3, 150 mM NaCl, 10 μM ThT) at 37 °C with shaking at 700 rpm. Insulin monomers were freshly prepared in the same buffer at pH 3 immediately prior to incubation. TEM Images of reduced insulin without lipids (c), with GM3 (d) and with GD3 (e). Indicated scale bars are 100 nm.

### 3.6. Insulin-ganglioside aggregates seed insulin aggregation

To investigate whether ganglioside-modified aggregates can act as nucleating seeds for insulin amyloid formation, ThT fluorescence assays were performed in the presence of 12.5%, 25% and 37.5% (molar percentage) of preformed insulin-ganglioside aggregates as seeds with 80 μM freshly prepared insulin monomers. The preformed insulin-ganglioside aggregates were centrifuged to separate the insulin-ganglioside species from the supernatant and these separated species (in the pellet) were used as seeds for aggregating insulin monomers. In contrast to ∼34-hour lag time observed for the insulin-alone control, the samples with preformed seeds showed accelerated aggregation kinetics, characterized by a significantly shortened lag time and faster rise in ThT fluorescence intensity (Fig. 10). The seeding effect was dose dependent, with the most pronounced accelerating observed at 37.5% seed concentration. Importantly, control experiments confirmed that the seed alone (in the absence of fresh insulin) exhibited negligible ThT fluorescence, ruling out the possibility that the signal originated from the seeds themselves. Structural analysis through circular dichroism revealed that both seeds form β-sheet rich structures, though the samples with insulin-GD3 seeds do contain some α-helical content as well (Fig. 10). In agreement, TEM micrographs showed well-defined fibrillar morphologies in the seeded samples, further confirming the ganglioside-modified aggregates retain the ability to nucleate amyloid fibril formation (Fig. 10). These results also demonstrate that insulin-ganglioside aggregates not only alter the aggregation pathway but also retain seeding capacity, suggesting a mechanism by which such species may promote further insulin amyloid propagation under physiological or pathological conditions.

**Figure 10.**
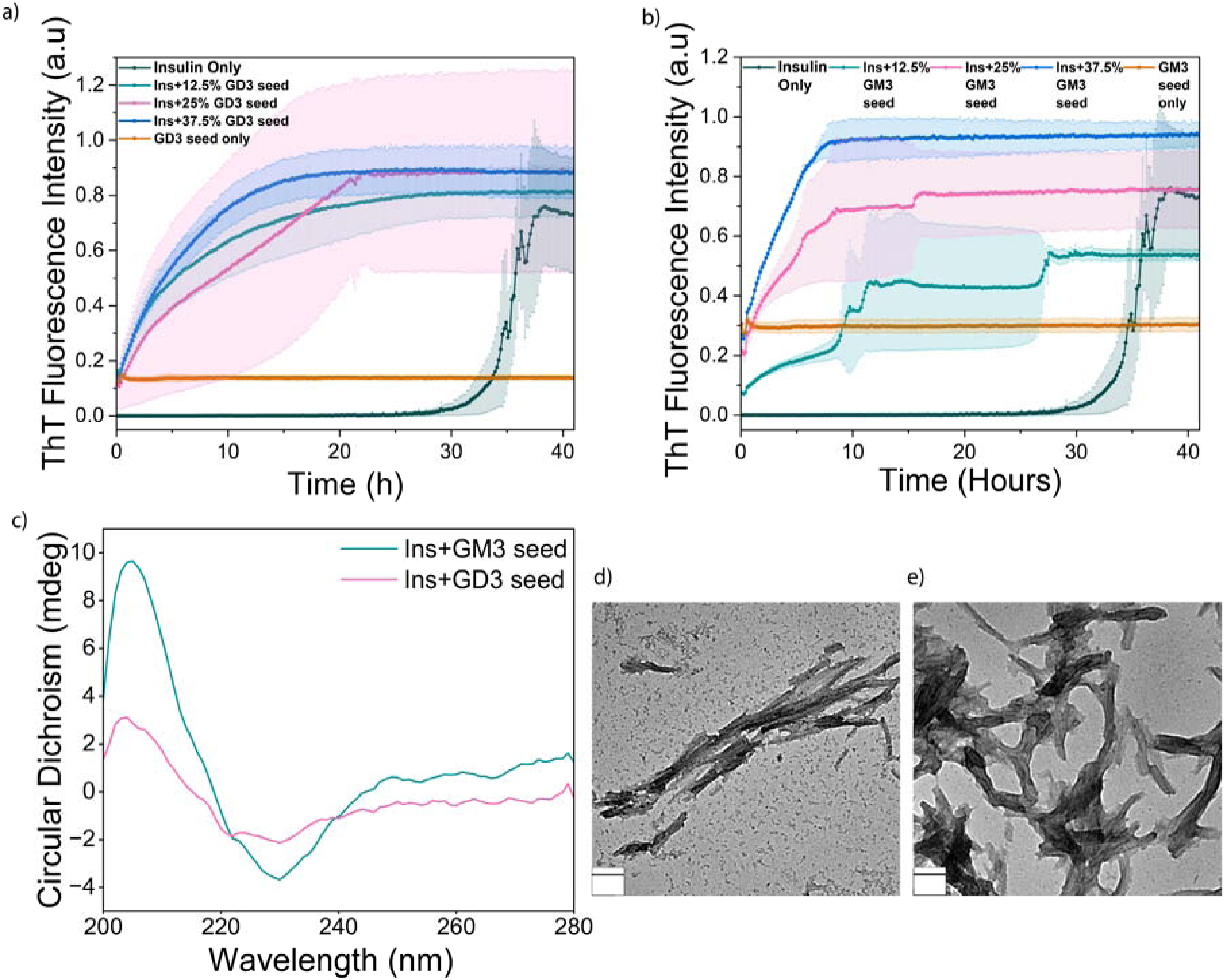
ThT Fluorescence kinetics of 80 μM insulin in the absence of seed (black) and in the presence of 12.5% (green), 25% (pink), and 37.5% (blue) (molar percentage) preformed (a) GD3-insulin and (b) GM3-insulin aggregates. Samples containing seeds alone without fresh insulin (dark green). (C) CD spectra of insulin aggregated in the presence of insulin-GD3 species (pink) and insulin-GM3 (green) seeds, demonstrating b-sheet-rich secondary structure. (d,e) TEM images of fibrils formed in the presence of (d) insulin-GD3 seeds and (e) insulin-GM3 seeds, confirming the presence of fibrillar morphologies. Indicated scale bars are 100 nm.

## 4. Discussion

Kinetic analysis of insulin aggregation in the presence of gangliosides GD3 and GM3 revealed a pronounced, concentration-dependent increase of the aggregation process, as shown by a marked reduction in lag time and t□/□ values obtained from the sigmoidal fitting of the ThT fluorescence curves. The 34 hours lag time for insulin control, which is consistent with slow nucleation, was reduced to around 9 hours in the presence of 5× concentration of both GD3 and GM3. A possible mechanism underlying the accelerated aggregation can be ganglioside micelles acting as conformational scaffolds, anchoring amyloidogenic proteins to membrane-like interfaces and accelerating structural reorganization. Similar to findings with Aβ, where GM1 micelles facilitated rapid β-sheet transition through peptide anchoring at the lipid sugar interface [68,69], our results suggest that GD3 and GM3 micelles bind insulin, acting as a scaffold and accelerating its aggregation. Another study from this same group found that the glycan cluster on the micelle surface of gangliosides stabilizes specific peptide folds by creating a hydrophobic/hydrophilic interface, which promotes ordered membrane interacting conformation, accelerating the aggregation process [70]. This behavior is also consistent with observations by Choo-Smith et al, where ganglioside vesicles seeded Aβ fibrillogenesis, which highlights their role in enhancing aggregation kinetics through membrane-mediated clustering [71]. The LVEALYL segment (residues B11–B17) of the insulin B-chain forms the highly ordered steric zipper spine of canonical insulin fibrils, where parallel β-strands assemble into tightly packed β-sheets [72,73]. The ability of GD3 and GM3 to shorten the lag phase may reflect their capacity to scaffold insulin monomers in conformations that partially expose or rearrange this critical segment, thereby facilitating early nucleation-like events. However, ganglioside binding may also trap LVEALYL-containing intermediates in non-productive conformations, preventing proper steric zipper assembly and ultimately diverting aggregation toward the off-pathway oligomeric species observed in our study. Notably, GD3 exhibited a stronger effect than GM3 at the intermediate concentrations of 1.5× and 2.5×, with GD3 exhibiting a lower lag time (around 14 hours) compared to GM3 at those concentrations (around 20 hours), suggesting that GD3 may more effectively promote nucleation-like events. This enhanced effect may stem from the higher negative charge (net charge -2) present on GD3 head group, which leads to stronger electrostatic interactions with the positively charged monomer of insulin at low pH, leading to earlier nucleation events. This is consistent with previous reports showing that higher negative charge on cardiolipin (net charge -2) leads to a shorter lag time in insulin aggregation, compared to phosphatidylglycerol with net charge -1 and control insulin without lipids [30]. These observations are further substantiated by the behavior observed in NMR spectra of insulin at pH 3.0. When insulin in no-salt conditions is exposed to GM3 micelles, immediate precipitation was observed with a loss in signal of ∼30% and 100% at 1× and 5× GM3, respectively. This effect was reduced by the presence of salt (100 mM NaCl), where only ∼17% signal was lost at 1× GM3. Notably, ∼80% of signal was lost due to rapid precipitation for 1× GD3 even in the presence of 100 mM NaCl. This is likely an effect stemming from the higher negative charge present on the surface of GD3 micelles. The ability of GD3 and GM3 to shorten the lag phase of insulin aggregation and the rapid, immediate aggregation behavior observed in NMR spectra points towards the role of electrostatic interaction steering in non-specific recruitment of the insulin molecule to the surface of the micelle at the early stages of the aggregation pathway.

The differential impacts of GD3 and GM3 may have also stemmed from the differences in the sialic acid content in GM3 and GD3 as well. Sialic acid plays a critical role in ganglioside–peptide interactions, as evidenced by studies showing that native GM1 induces structural changes in Aβ, while asialo-GM1, which lacks sialic acid, does not elicit such conformational transitions [71,74,75]. Kakio et al. also found that in accelerated fibrillogenesis of Aβ via binding to gangliosides, the number of binding sites on the gangliosides are roughly proportional to the number of sialic acid residues in the lipids [76]. The molecular dynamics simulations provided complementary atomistic support for this hypothesis, showing that the insulin monomers preferentially localize at the ganglioside bilayer interface. The potential of mean force profiles indicated that insulin favors surface absorption at ∼4–4.5 nm distance from the bilayer center, with binding free energies of 2–4 kJ/mol. This moderate but favorable interaction likely stabilizes the membrane-bound intermediate states necessary for the early nucleation-like events. Furthermore, protonation under acidic pH increased lipid contact frequency, especially for positively charged B-chain residues (Phe1-B, Val2-B, Hsp5-B, Hsp10-B), which enhanced the electrostatic interactions between insulin and the negatively charged ganglioside headgroups. Our results, along with previous literature, highlight the capacity of gangliosides to modulate the early kinetic events of insulin amyloidogenesis based on their charge density and colloidal state.

Our structural characterization of insulin-ganglioside aggregates using TEM, SAXS, FTIR and CD reveals that GM3 and GD3 not only accelerate insulin aggregation but also shift it away from the classical amyloid formation pathway and lead towards morphologically and structurally distinct aggregates. While insulin alone formed long, unbranched β-sheet rich fibrils consistent with canonical amyloid morphology, the presence of gangliosides at all tested concentrations led to the formation of non-fibrillar, globular or beaded aggregates, which lacked the uniformity and length of typical amyloid fibrils. These morphologies imply the formation of off-pathway oligomers, which are assemblies that deviate from the nucleation-dependent fibrillization pathway and do not progress into mature fibrils. Markedly, GD3 at 1.5× and 5× concentration promoted aggregates with remarkably high β-sheet content as seen in the FTIR deconvolution despite the absence of fibrillar structure, suggesting the formation of β-sheet rich oligomeric species. This aligns with multiple studies reporting β-sheet-rich oligomeric intermediates, including a study by Sangwan et al. where they showed that Aβ fragments form antiparallel, out of-register β-sheet oligomers with no fibrillar morphology [77]. Similarly, some inositol stereoisomers also stabilized β-structured Aβ42, confirmed by CD, without permitting fibril elongation [78]. In contrast, GM3 induced aggregates with a concentration-dependent increase in α-helical content. This suggests that GM3 may stabilize native-like or helical intermediates, stopping the complete structural transition into long β-sheet fibrils. The appearance of shoulders at 1655 cm^-1^ in FTIR spectra and increased signal at 208 and 222 nm in CD spectra further supports this interpretation. This is consistent with studies showing that GM1 and GM2 micelles induce α-helical conformation in Aβ40. NMR based torsion angle analysis also confirms that specific segments of Aβ adopt discontinuous α-helices upon binding to ganglioside. This indicates that similar helical intermediates may form during insulin interaction with GM3 and GD3 [68,74]. These findings are consistent with reports that lipid environments can trap proteins in non-fibrillar states [79], either by stabilizing the early oligomeric intermediates or by disrupting the β-sheet alignment. Liu et al. resolved macrocyclic β-sheet tetramers which formed non-fibrillar, trapped states due to geometric constraints, suggesting that off-pathway oligomers can be both energetically and sterically stabilized [79]. Similarly, Cerf et al. provided evidence of Aβ oligomers adopting antiparallel β-sheet structure persisting as stable, off-pathway conformations [80]. Molecular simulations of ATTR (105-115) peptide assemblies reinforce this model [81]. In this study, the formation of β-sheet-rich oligomers was also observed, with hydrophobic residues driving the formation of β-barrel-like structures, which acted as stable intermediates or even distinct thermodynamic endpoint on an alternative aggregation pathway. SAXS data provides further evidence that GM3 and GD3 modulate the structural organization of insulin aggregates. The emergence of Bragg peaks and lattice-like packing in high-concentration ganglioside samples support the presence of ordered, non-fibrillar macro-structures. This structural order, combined with the globular morphology seen in TEM and reduced β-sheet signal in FTIR/CD, reinforces the idea that GM3 and GD3 differentially remodel insulin aggregation pathways, promoting off-pathway oligomeric species formation. These results are further supported by the simulation data, which revealed that GM3 and GD3 differentially engage specific insulin residues at the bilayer surface. Under acidic conditions (at pH 3), ganglioside binding was dominated by defined polar contacts involving B-chain helical residues. This spatially restricted interaction could stabilize native-like or partially folded conformers, preventing β-sheet formation and fibril propagation. The observed differences in area per lipid (0.849 nm² for GD3 vs. 0.678 nm² for GM3) may also affect bilayer fluidity and interaction surface geometry, which could influence the differential aggregate morphology. This interpretation is consistent with the observation that ganglioside-induced aggregates fail to progress into mature fibrils; ganglioside binding may stabilize β-sheet-rich intermediates in which the LVEALYL segment is partially engaged but unable to form the fully interdigitated steric zipper required for fibril elongation.

The secondary structure content derived from deconvolution of both CD and FTIR spectra shows differences in the absolute percentage of α-helical and β-sheet structures. While CD and FTIR are utilized as complementary techniques, the physical principles underlying these two techniques are fundamentally different. CD spectroscopy probes the differential absorption of left- and right-circularly polarized light, which makes it particularly sensitive to α-helical content in soluble form but that also makes it less responsive to rigid, surface-constrained or densely packed formations [82,83]. In contrast, FTIR spectroscopy probes the backbone hydrogen bonding through the amide I bond and picks up both soluble and insoluble protein content, which includes structures immobilized or constrained within aggregates [84,85].Additionally, researchers have shown before that protein structures adsorbed onto oil-water interface produce a significant amount of light-scattering, which interferes with measuring secondary structure of adsorbed protein by CD [86]. In the insulin-ganglioside system being studied, the observed difference possibly arises from the presence of ganglioside-bound aggregates that are partially rigid and adsorbed onto the lipid surface, leading to suppression of CD signal. This is supported by the NMR results, which showed complete or partial signal loss in the presence of GD3 and GM3, indicating large, precipitating aggregates. Molecular simulation also showed that insulin stably binds to ganglioside surface, supporting the fact that the structural transitions occur in membrane bound, restricted context. These findings support the notion that FTIR captures the secondary structure of ganglioside-bound insulin aggregates, while CD is unable to represent these structures due to immobilization and the presence of large ganglioside micelles. Interestingly, when Oberg et al. explored the complementary behavior of CD and IR spectra, they also noted that if the α-helical content from CD or FTIR is significantly lower than the other, the larger amount might be closer to reality [83,87].

The distinct secondary structure and morphology of the ganglioside-insulin aggregates had important functional implications as well. While the insulin-only aggregates were highly toxic to the NIH3T3 cells, the aggregates formed with GD3 and GM3 demonstrated significantly reduced toxicity, particularly after the removal of unbound gangliosides via filtration. It is possible that gangliosides remain stably associated with insulin aggregates at the end point of the reaction, due to their amphiphilic nature and strong affinity for protein surfaces, including potential electrostatic interactions with insulin. This behavior is consistent with previous observations of ganglioside binding to Aβ [88]. While β-sheet rich oligomers have been reported to be toxic, the insulin-ganglioside oligomers observed in our study appear to diverge from these canonical toxic species. This divergence may be attributed to surface shielding effects conferred by the gangliosides. These bound ganglioside molecules could act as structural modulators by masking the hydrophobic or membrane active regions of insulin aggregates and thus reduce their capacity to interact with cellular membranes, a key mechanism underlying amyloid-induced toxicity [89–93]. Previous literature has noted that not all β-sheet-rich oligomers are cytotoxic. Instead, toxicity depends on the membrane interaction potential and exposure of the hydrophobic patches. In our system, GD3 and GM3 may sequester insulin into compact β-rich conformation with a persistent ganglioside association, leading to structurally inert and membrane-inactive species, thereby mitigating their toxicity [94]. The observation that non-filtered samples retained some toxicity, along with mild toxicity observed for lipid only controls, suggests that the protective effect of the insulin-ganglioside aggregates does not stem from the presence of ganglioside in the sample. Instead, it likely arises from their association with insulin and the resulting structural modifications induced by the GD3 and GM3.

The aggregation behavior of reduced insulin differed markedly from that of native insulin. The ThT fluorescence assays showed a complete absence of fibril formation, regardless of the presence of gangliosides, and TEM images confirmed the presence of amorphous, non-fibrillar aggregates. These findings indicate that GD3 and GM3 do not influence the aggregation of reduced insulin. This suggests that gangliosides alone are not sufficient to nucleate ordered aggregation in the absence of disulfide-stabilized insulin conformations. Instead, the reduced system favors disordered, ThT-inactive aggregates regardless of the lipid environment. Notably, however, the GD3 and GM3-induced insulin aggregates were capable of self-seeding insulin aggregation when incubated with native monomeric insulin, shortening the lag time significantly. This demonstrates their capacity to act as template for amyloid growth and highlights a distinct difference in how gangliosides modulate aggregation depending on the structural state of the insulin monomer. This was further supported by CD spectra and TEM images respectively showing β-sheet structure and fibrillar morphology. These findings suggest that, despite their non-fibrillar and globular morphology, insulin-ganglioside aggregates retain surface β-structure motifs or oligomeric templates that are capable of secondary nucleation and elongation. This is in agreement with a previous report on insulin’s self-seeding capability its own amyloid aggregation [95]. Furthermore, it is consistent with reports demonstrating that non-canonical, prefibrillar oligomers can act as nucleation competent surfaces [76,80,96].

While our findings provide compelling evidence that gangliosides GM3 and GD3 modulate insulin aggregation kinetics, structure, and cytotoxicity in vitro, several important questions remain for future investigation. Testing the effect of gangliosides and insulin aggregates in animal models, such as diabetic mice, could help determine the physiological relevance of these aggregates and their potential role in amyloid formation in vivo. Additionally, reconstituting the gangliosides into lipid bilayers or membrane rafts could also help determine how membrane presentation and microdomain organization influence aggregation, since gangliosides naturally reside in membrane environments. Molecular Dynamics simulations could further reveal atomic-level insights into the ganglioside-insulin interactions, potentially elucidating the mechanism underlying structural modulation. Finally, investigating whether other disulfide-rich or α-helical protein respond similarly to gangliosides could reveal broad principles governing the modulatory effects of gangliosides on protein aggregation.

## 5. Acknowledgements

We thank Dr. Samuel McCalpin, Dr. Stephen Arce, Emily Thiel, Dr. Jamel Ali, and Dr. Navneet Kaur for valuable discussions and technical assistance. SAXS/WAXS measurements were performed under the user program of the Center for Nanophase Materials Sciences (CNMS), a U.S. Department of Energy Office of Science User Facility operated by Oak Ridge National Laboratory. We also acknowledge the use of instrumentation and core facilities at the Biological Science Imaging Resource (BSIR), the Institute of Molecular Biophysics (IMB), and the High-Performance Materials Institute (HPMI). We are especially grateful to Peter Randolph at IMB and Anthony Warrington at BSIR for the technical support. Computational resources were provided by the Florida State University Research Computing Center.

## 6. Author Contributions

**Nazifa Tasnim Ahmad**: Investigation, Methodology, Data curation, Formal analysis, Visualization, Writing – original draft, Writing – review & editing. **Jhinuk Saha**: Investigation, Methodology, Data curation, Writing – review & editing. **Yimin Mao**: Investigation, Writing – review & editing. **Robert Silvers**: Investigation, funding acquisition Writing – review & editing. **Zaid Abulaban**: Investigation. **Joshua Mysona**: Investigation, Writing – review & editing. **Ayyalusamy Ramamoorthy**: Conceptualization, Methodology, Supervision, Funding acquisition, Project administration, Writing – review & editing.

