## Supporting Information for "Gangliosides GM3 And GD3 Modulate Insulin Aggregation Pathways and Reduce Cytotoxicity Through Structural Remodeling"

### Supplementary Information

(a)

(b)

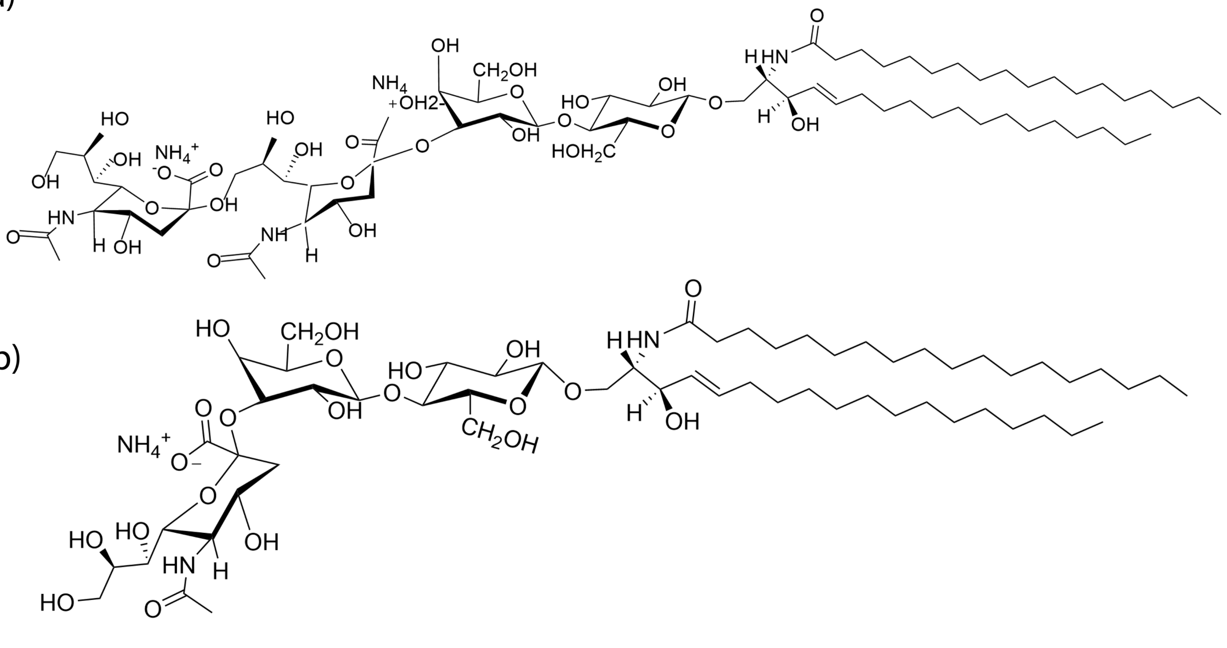

**Figure S1**. Molecular structures of (a) GD3 and (b) GM3 lipids used in this study.

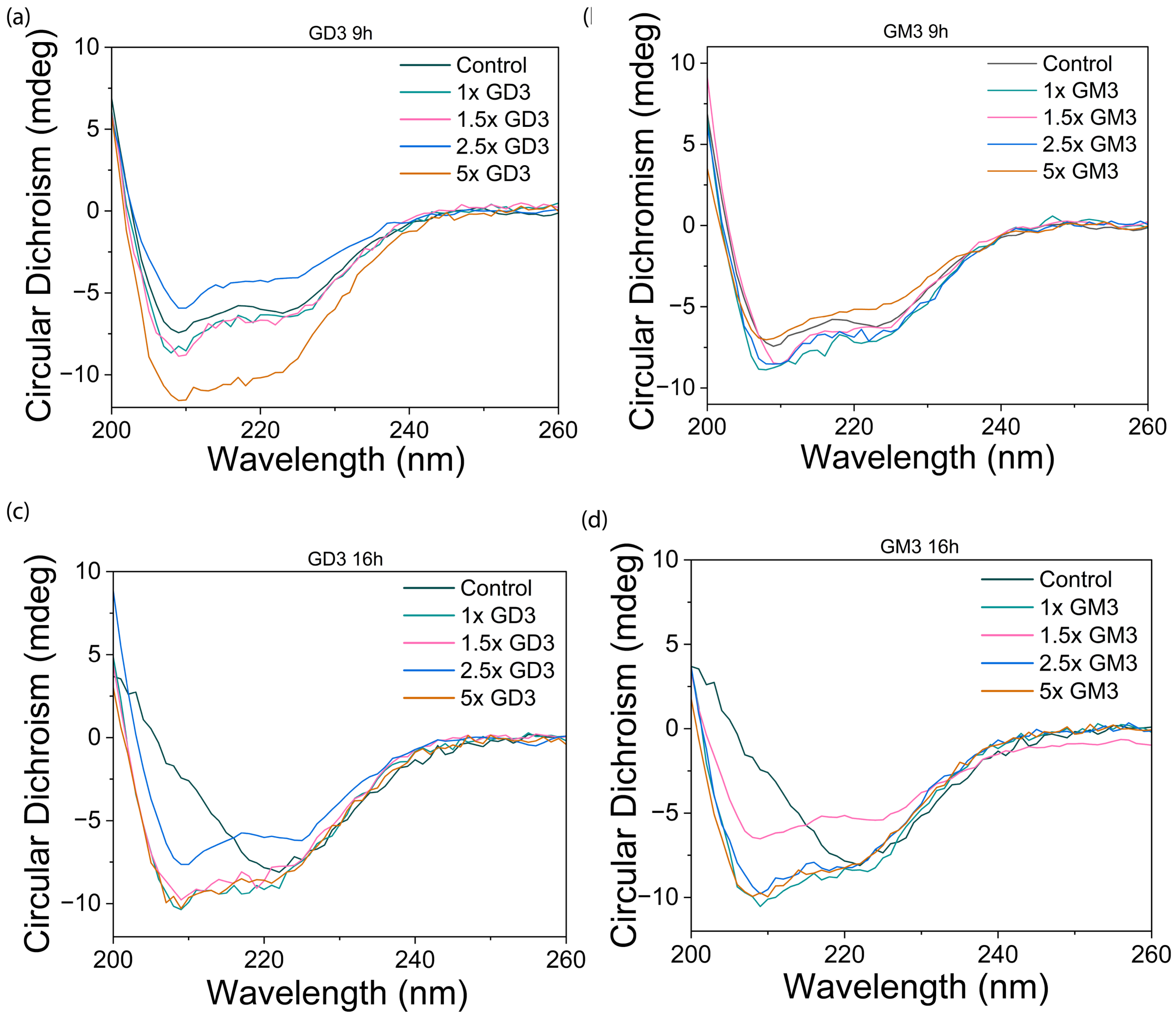


**Figure S2**. CD spectra of 80 μM insulin in a lipid free environment (dark green) and in the presence of increasing concentrations of gangliosides, GD3 (a and c) and GM3 (b and d): 1× (green), 1.5× (pink), 2.5× (blue), and 5× (yellow). (a) GD3 and b) GM3 after 9(a,b) and 16(c,d) hours of incubation.

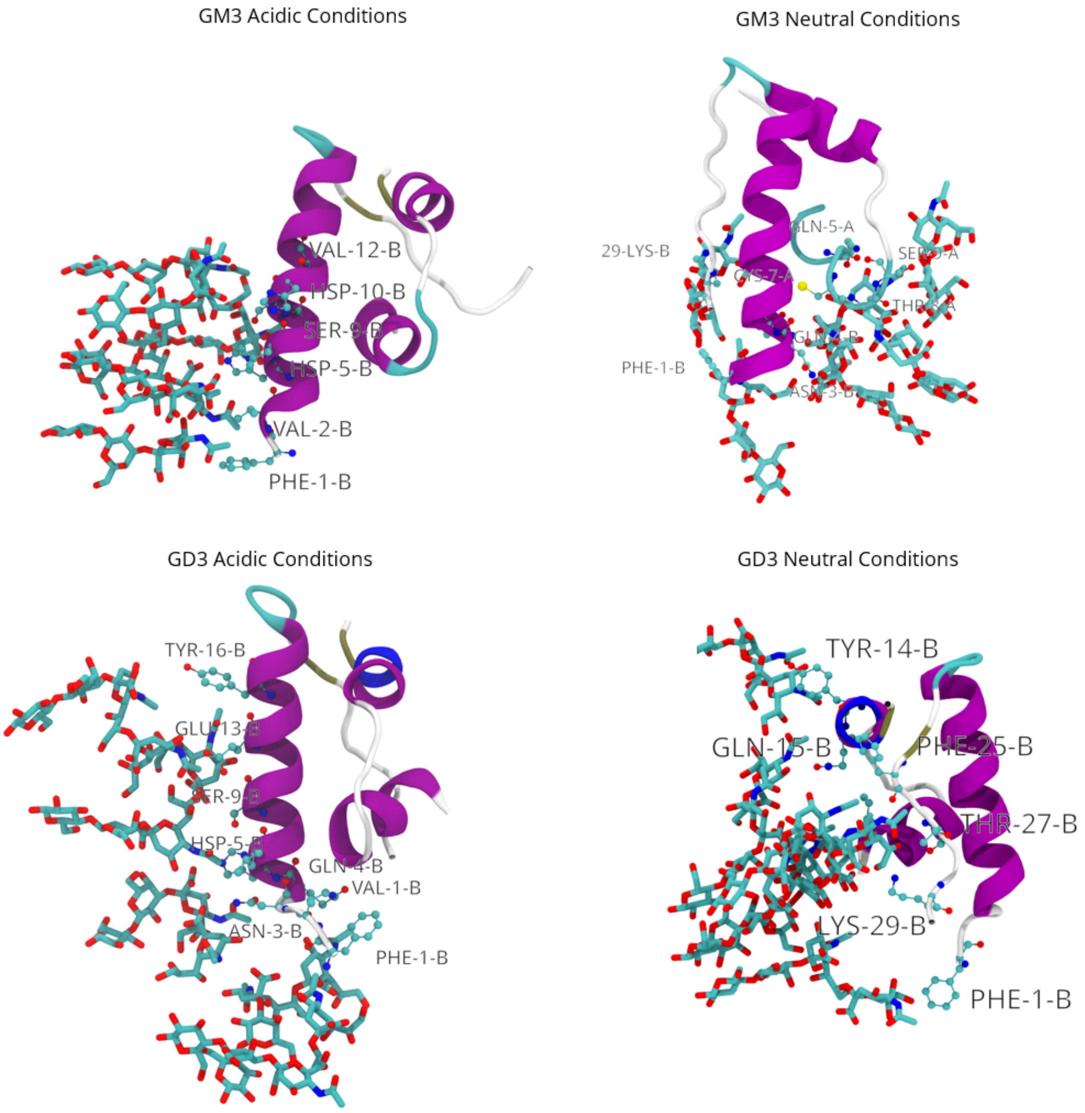


**Figure S3**. Detailed representative simulation snapshots of the insulin molecule on the bilayer surface. Polar GM3/GD3 head groups interacting with the molecular are drawn as well as the relevant protein residues.

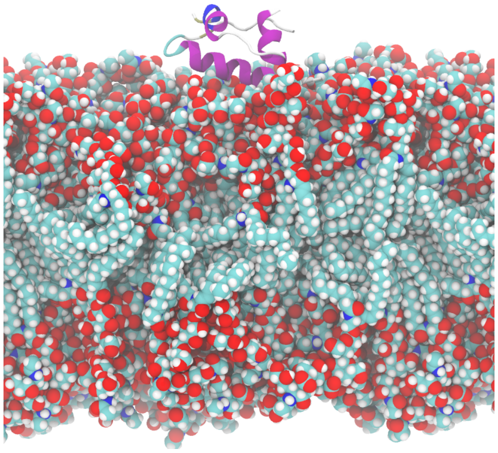

**Figure S4**. Insulin molecule interacting with the GD3 bilayer surface.

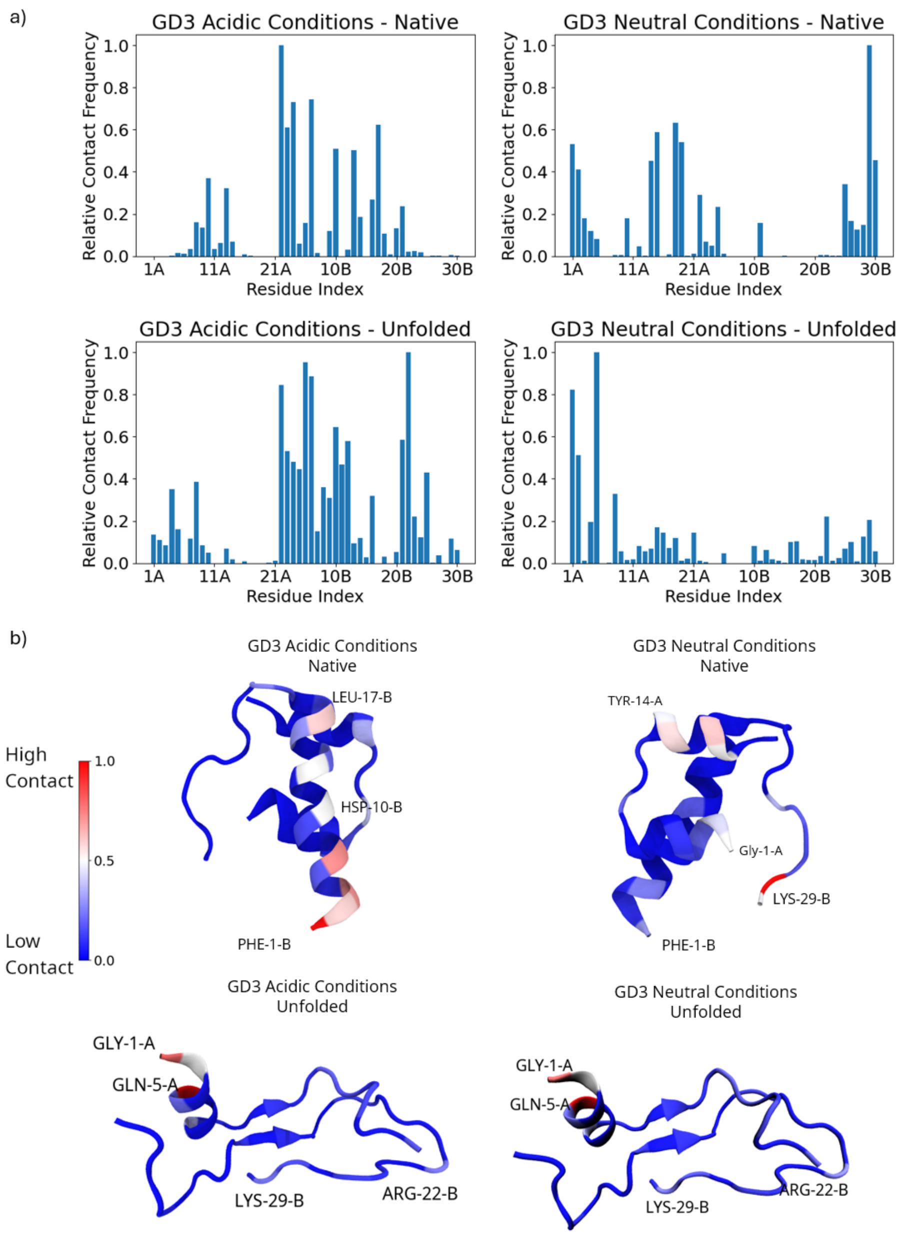


**Figure S5**. (a) Plot of contact frequency between each residue of the insulin and GD3 lipid bilayer normalized to the most frequent in contact residue. (b) Insulin monomers with residues are colored by overall contact frequency: red as the most frequent and blue as the least frequent.

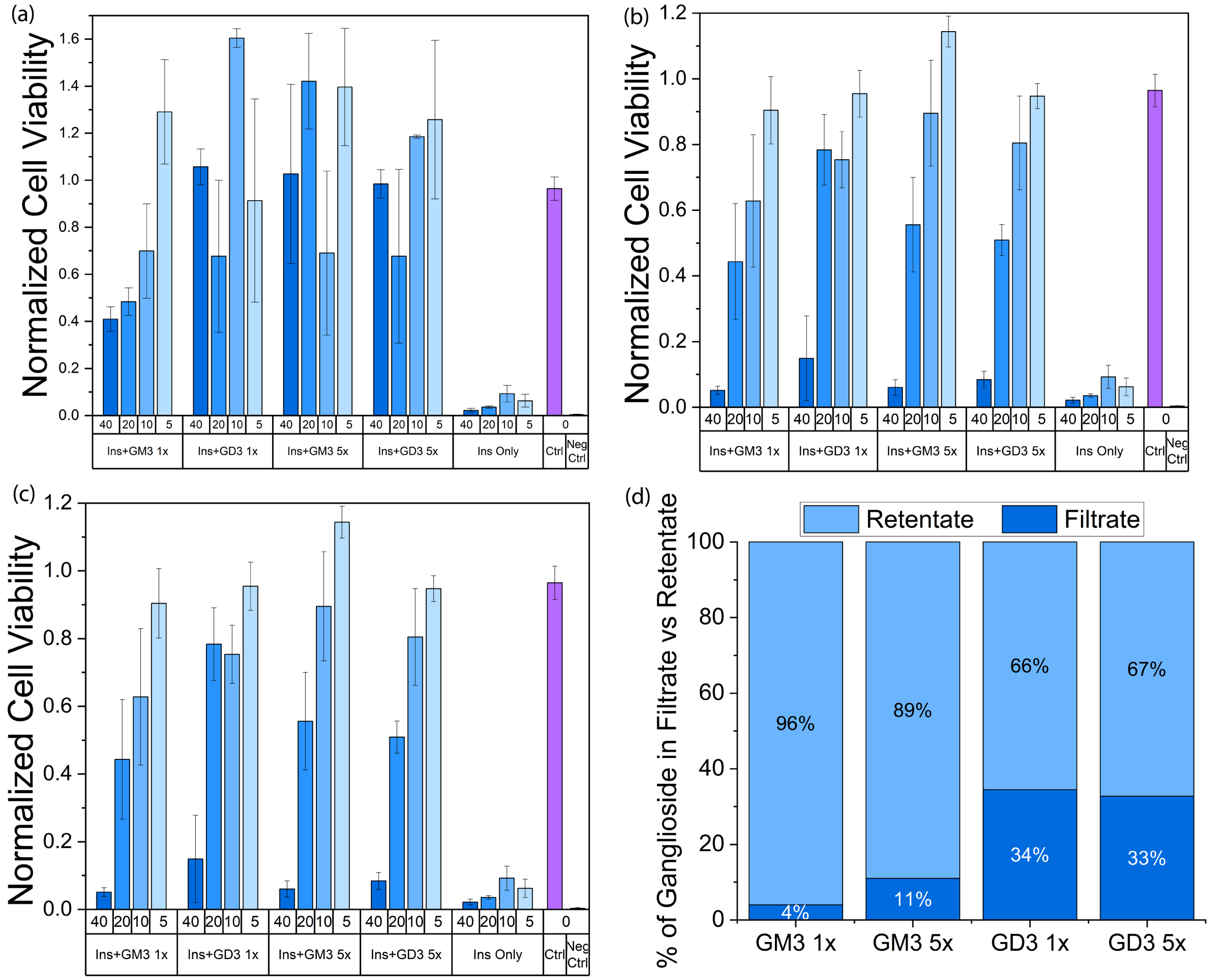


**Figure S6**. Viability of NIH3T3 cells incubated with (a) filtered, (b) non-filtered insulin-ganglioside aggregates and (c) only gangliosides. Cell viability was assessed using the XTT cell viability assay for NIH3T3 cells after 24 hours of incubation with the indicated samples. Non-filtered samples contained both aggregates and unbound gangliosides, while filtered samples contained only aggregates. (c) Percentage distribution of ganglioside content in the filtrate and retentate fractions of the insulin-ganglioside samples after filtration, showing the relative distribution of GM3 and GD3 at 1× and 5× molar excess conditions.

| *Sample* | $F_{MAX}$ | *Slope* | $T_{1/2}$ *(h)* | $T_{LAG}$ *(h)* |
|---|---|---|---|---|
| *GD3 1×* | 0.5 | 0.7 | 19.9 | 18.4 |
| *GD3 1.5×* | 0.4 | 1.7 | 15.7 | 15.1 |
| *GD3 2.5×* | 0.4 | 0.7 | 16.2 | 14.7 |
| *GD3 5×* | 0.8 | 0.8 | 10.7 | 9.4 |
| *GM3 1×* | 0.7 | 1.1 | 26.1 | 25.2 |
| *GM3 1.5×* | 0.7 | 1.4 | 20.3 | 19.6 |
| *GM3 2.5×* | 0.7 | 1.1 | 22.5 | 21.6 |
| *GM3 5×* | 0.7 | 0.9 | 11.5 | 10.3 |
| *Only Insulin* | 0.6 | 1.0 | 35.1 | 34.1 |

**Table S1**. Fitted kinetic parameters from sigmoidal analysis of ThT fluorescence curves for insulin aggregation. Parameters include lag time (tlag), half-time ($t_1/_2$), maximum fluorescence intensity (Fmax), and apparent elongation rate (slope).

| **Sample** | **GD3 1×** | **GD3 1.5×** | **GD3 2.5×** | **GD3 5×** | **GM3 1×** | **GM3 1.5×** | **GM3 2.5×** | **GM3 5×** | **Only Insulin** |
|---|---|---|---|---|---|---|---|---|---|
| Alpha helix | 61.7 | 3.6 | 50.4 | 11.8 | 0.0 | 18.1 | 28.2 | 41.6 | 3.8 |
| Antiparallel Beta sheet | 1.2 | 0.4 | 0.4 | 0.0 | 0.7 | 0.0 | 0.6 | 0.0 | 0.3 |
| Beta sheet | 31.3 | 87.5 | 43.4 | 73.9 | 41.3 | 43.6 | 61.3 | 43.6 | 47.6 |
| Turn | 5.8 | 8.6 | 5.8 | 12.7 | 58.0 | 35.6 | 8.9 | 14.7 | 48.3 |
| Random coil | 0.0 | 0.0 | 0.0 | 1.6 | 0.0 | 2.7 | 1.0 | 0.0 | 0.0 |

**Table S2.** Estimated secondary structure composition of insulin-ganglioside aggregates based on FTIR analysis in the Amide I region (1600-1700 $cm^{-1}$). Samples contained 80 µM insulin with 1×, 1.5×, 2.5× and 5× molar equivalents of GD3 and GM3, incubated in 10 mM pH3 sodium phosphate buffer containing 150 mM NaCl.

| Samples | GD3 1× | GD3 1.5× | GD3 2.5× | GD3 5× | GM3 1× | GM3 1.5× | GM3 2.5× | GM3 5× | Only Insulin |
|---|---|---|---|---|---|---|---|---|---|
| Helix | 5.8 | 4.5 | 5.9 | 4.1 | 4.8 | 2.7 | 5.1 | 7.4 | 0 |
| Antiparallel | 31.3 | 32.6 | 34.3 | 32.5 | 33.6 | 36.1 | 34.8 | 31 | 34.6 |
| Parallel | 0.0 | 0 | 0 | 0 | 0 | 0 | 0 | 0 | 0 |
| Turn | 14.9 | 15.2 | 13.1 | 14.7 | 15.6 | 14.7 | 14.9 | 14.8 | 20.5 |
| Others | 48.0 | 47.7 | 46.8 | 48.6 | 46 | 46.5 | 45.2 | 46.7 | 44.9 |

**Table S3**. Estimated secondary structure composition of insulin-ganglioside aggregates based on CD spectra. Samples contained 80 µM insulin with 1×, 1.5×, 2.5× and 5× molar equivalents of GD3 and GM3, prepared in 10 mM pH3 sodium phosphate buffer containing 150 mM NaCl.

**SV1.** Representative videos of molecular dynamics trajectories illustrating insulin rotational dynamics when bound to ganglioside-containing lipid bilayers under different lipid compositions and pH conditions: insulin bound to a (a) GM3-containing bilayer under acidic conditions; (b) GM3-containing bilayer under neutral conditions; (c) GD3-containing bilayer under acidic conditions; (d) GD3-containing bilayer under neutral conditions.
